# Homologous Pairs of Low and High Temperature Originating Proteins Spanning the Known Prokaryotic Universe

**DOI:** 10.1101/2023.08.24.554664

**Authors:** Evan Komp, Humood Alanzi, Ryan Francis, Chau Vuong, Logan Roberts, Amin Mossallenejad, David A. C. Beck

## Abstract

Stability of proteins at high temperature has been a topic of interest for many years, as this attribute is favourable for applications ranging from therapeutics to industrial chemical manufacturing. Our current understanding and methods for designing high-temperature stability into target proteins are inadequate. To drive innovation in this space, we have curated a large dataset, learn2thermDB, of protein-temperature examples, totalling 24 million instances, and paired proteins across temperatures based on homology, yielding 69 million protein pairs - orders of magnitude larger than the current largest. This important step of pairing allows for study of high-temperature stability in a sequence-dependent manner in the big data era. The data pipeline is parameterized and open, allowing it to be tuned by downstream users. We further show that the data contains signal for deep learning. This data offers a new doorway towards thermal stability design models.

## Background & Summary

High-temperature proteins (HTPs) find their way into wide-ranging applications spanning from drugs and diagnostics in human health to catalysing reactions in industrial processes and bioremediation.^1–5^ The extraordinary adaptability of these proteins enables them to maintain functionality even in extreme conditions— a feat that unfortunately remains a formidable challenge in protein design and engineering.^6–8^ One key determinant of protein adaptability is thermal stability, governed by the Gibbs free energy difference between the folded and native states. This energy balance, crucial for proteins to maintain their functional conformation, remarkably holds across various organisms and environments.^9,10^ Even so, minor alterations in protein sequences can have profound impacts on their intricate structures, and consequently, on their stability and folding capacity.^11^ Despite these challenges, life finds a way: thermophiles, which rely on HTPs, can thrive at extreme temperatures.^12–15^

Current methods that aim to produce proteins functional at higher temperatures, such as directed evolution or rational design, unfortunately, provide no guarantee of success and require substantial effort for each new protein of interest.^16–24^ Recent advances in fully deep learned designers have shown impressive results and generating sequences that reliably fold to a target structure, particularly at ambient temperatures.^25–28^ While new and improved models are continuing to be developed, they do not invariably produce designs that retain tertiary structure and activity at high temperatures.^29–31^ Strategies which are based on substructures, energy functions, or patterns learned from protein structures represented in the PDB, are limited considering that the majority of proteins are non-thermophilic: only 5% of proteins from the top 25 most populous source organisms are thermophilic.^32–35^ The temperature-dependent nature of enthalpic and entropic forces in the protein means that stability at ambient temperature does not necessarily translate to high-temperature stability.^36,37^ Learning from high temperature proteins in a targeted manner may further improve reliability of novel methods. Consequently, researchers have been in search of rules employed by nature to produce HTPs for years.^14,38–42^ Table 1 presents a selection of datasets that have been utilised in this search for design principles.

**Table 1:** A selection of the largest publicly available datasets of proteins and their thermal stability.

| Name | Size Proteins | Size Prot. Pairs | Diversity | Label |
| --- | --- | --- | --- | --- |
| Meltome Atlas<sup>43</sup> | 48k | - | Diverse proteins, 13 organisms | T<sub>M</sub> |
| ProThermDB<sup>44</sup> | 32k | - | Diverse proteins and single point mutation | T<sub>M</sub> |
| FireProtDB<sup>45</sup> | 16k | - | Single point mutations | T<sub>M</sub> |
| Engqvist et al.<sup>46</sup> | 5.5 mil | - | Diverse enzymes, multiple examples across evolution | OGT |
| TemStaPro<sup>47</sup> | 2.5 mil | - | Diverse proteins, multiple examples across evolution | OGT |
| Hait et al.<sup>48</sup> | 700 | 1.6k | Diverse protein pairs | OGT |
| **This work (learn2therm database)** | **24 mil** | **69 mil** | **Diverse protein pairs, multiple examples across evolution** | OGT |

Existing datasets, apart from their size, suffer from a number of limitations. For example, single point mutation datasets such as FireProtDB allow for the study of stability independent from confounding factors like evolutionary drift, yet they offer limited variety and informational density when compared to the variety of known proteins.^37,45,49^ Larger datasets that label substantial portions of proteins using parameters such as melting temperature (T_M_) or the more noisy optimal growth temperature (OGT) of the host organism can be used to discern average trends.^50–53^ However, any thermal stability modes extracted in this way may not be universally applicable, as thermal stability is often specific to the protein fold itself.^42,43,54^ Therefore, patterns identified on average may even prove destabilising when applied to a new target.^41,55^ To account for this, thermal stability should be studied in a context dependent manner, where homologs are paired across temperature, with multiple examples across evolution. The largest dataset of protein pairs contains only 1.6k unique examples, and does not have redundancy across evolution.^48^

Recent advancements have shown that large deep learning models can effectively understand protein context in applications such as binding predictions, structure prediction, and backward design, among others.^30,56–61^ Each of these applications, however, is predicated on large (>10k) datasets. To translate these successes to context-dependent thermal stability design, the development of a large dataset of homologous protein pairs across different temperatures becomes imperative. This work introduces such a dataset: learn2thermDB. The dataset includes 69 million protein pairs of 250 amino acids or fewer derived from 4739 mesophilic organisms (with OGT < 40°C) and 289 thermophilic organisms (with OGT >40 °C, up to 98 °C) using homology search. We first paired mesophilic organisms to thermophilic ones by evolutionary distance, and then homologous proteins among those taxa pairs. Applying a stricter thermophilic threshold of 60 °C, used by the current largest dataset, our database still retains 9 million protein pairs, a four orders of magnitude increase. It is worth noting that individual proteins often participate in multiple pairs, providing the database with some redundancy across the evolutionary landscape. Moreover, even those organisms that did not demonstrate pairing still contribute to the dataset with their proteomes labelled by OGT, yielding 23 million proteins from mesophiles and 1 million from thermophiles (with a maximum OGT of 102 °C) for further study. The dataset’s size, in terms of the included organisms and proteins, is depicted in Table 2. The featured organisms span a broad range of prokaryotes across the taxonomic tree (Figure 1-left) and cover much of the known protein space (Figure 1-right). The current largest dataset of protein pairs across temperature from Hait et al. is also depicted for comparison in Figure 1-right. The distribution of OGT difference between organisms is shown in Figure 2-top, and the number of protein pairs as a function of OGT difference in Figure 2-bottom. We can see that the distribution is skewed towards similar temperatures, however we still retain millions of protein pairs with OGT difference >30 °C. The dataset as described in this manuscript is made available as-is on Figshare, please see the section “Data Descriptor” below.^62^

**Table 2:**
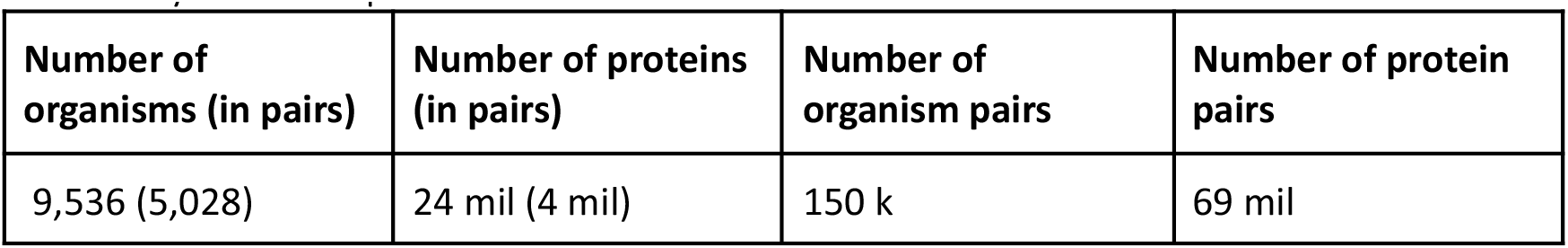
Number of various entities in the dataset all with temperature labels. Organism and protein pairs were identified via local alignment, thus not all taxa/proteins participate in pairing, and those that do may occur multiple times.

**Figure 1.**
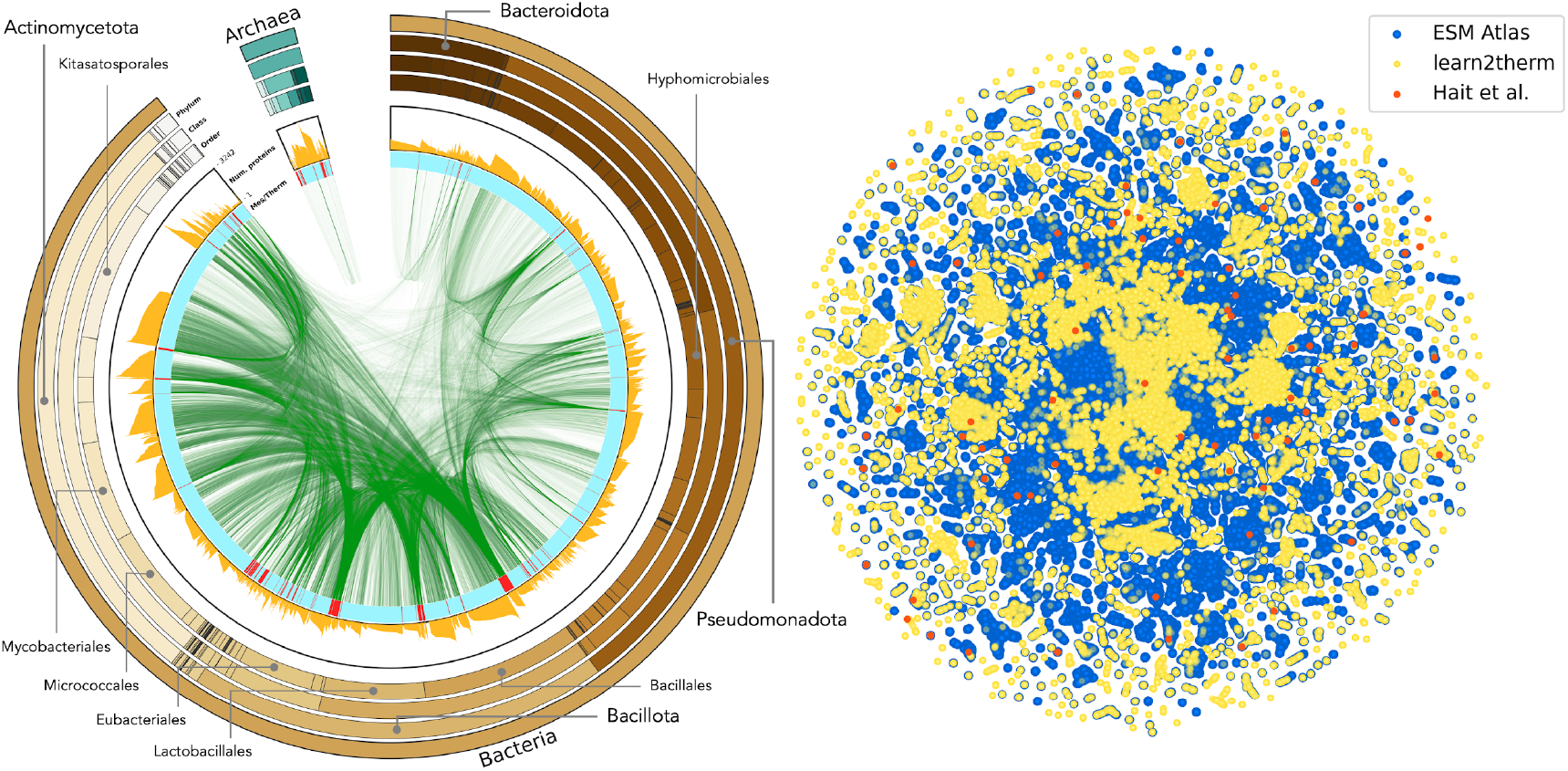
The taxonomic and protein space covered by the protein homologous pairs (N=69 mil) within the dataset. Left) NCBI taxonomic breakdown of the dataset.^63^ The outer ring depicts super kingdom, Phylum, Class, and Order, where the size of wedges indicates the number of organisms in the classification that contain at least one protein in a learn2therm protein pair. Highly populated Phylum and Class are labelled. The inner ring moving inward is a histogram of the number of proteins participating in pairs per organism followed by a colour mapping labelling organisms as mesophilic in blue and thermophilic in red. Central connections indicate taxa pairs contributing to protein pairs. Right) Two dimensional mapping (using t-SNE) of a sample protein space as determined by Evolutionary Scale Model (ESM) embeddings.^59,64^ In blue, a sample of data from the ESM Atlas with highest structural confidence. Note that this data contains eukaryotic proteins. In yellow, our proteins in pairs, and in orange, the current largest set of protein pairs across temperature. Size of samples conserves relative size of our proteins vs. the Atlas and reference dataset. For details of the mapping procedure, see Supplementary Information S8.

**Figure 2.**
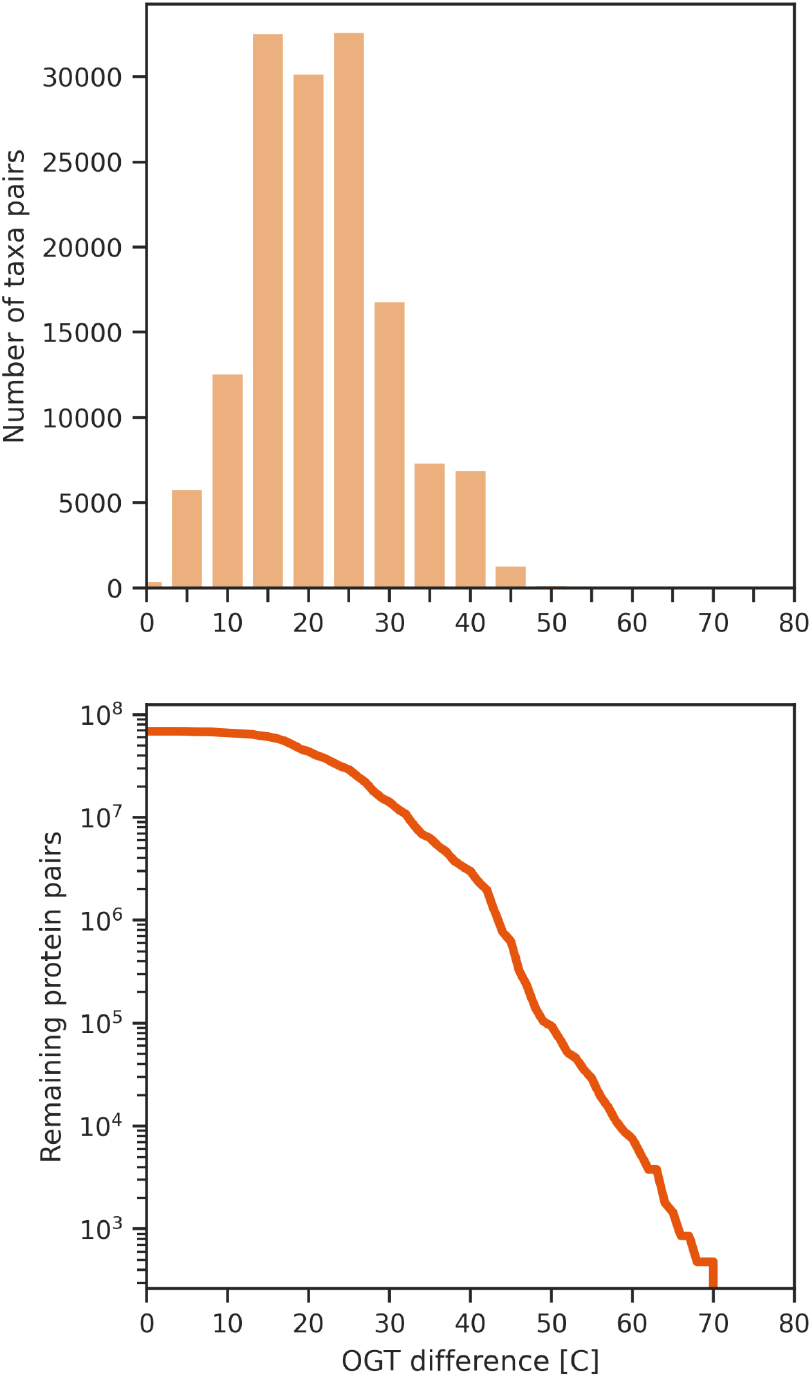
The data distribution as a function of difference in growth temperature between items. top) Histogram of difference in OGT between pairs of organisms. The data is skewed towards 0, but there are still many pairs with temperature differences >30 °C. bottom) Count of protein pairs remaining as minimum OGT difference is increased. We still retain 14 mil protein pairs with OGT difference >30 °C.

In addition to the static dataset provided, the entire pipeline used to produce the data is not only open source, but parameterized, such that an interested reader can reproduce or alter the produced dataset. Please see section “Usage Notes.” We believe that the carbon cost of compute efforts is as important as the result itself, thus our pipeline tracks approximate carbon cost.^65^ The cost of running the pipeline was estimated to be 11.5 kg carbon, according to the energy partition in Washington State, USA, and the compute total carbon cost of this research effort including development was estimated at about 70 kg.

## Methods

Our process of identifying homologous protein pairs across temperature is discussed in the following sections. The pipeline is depicted in Figure 3.

**Figure 3.**
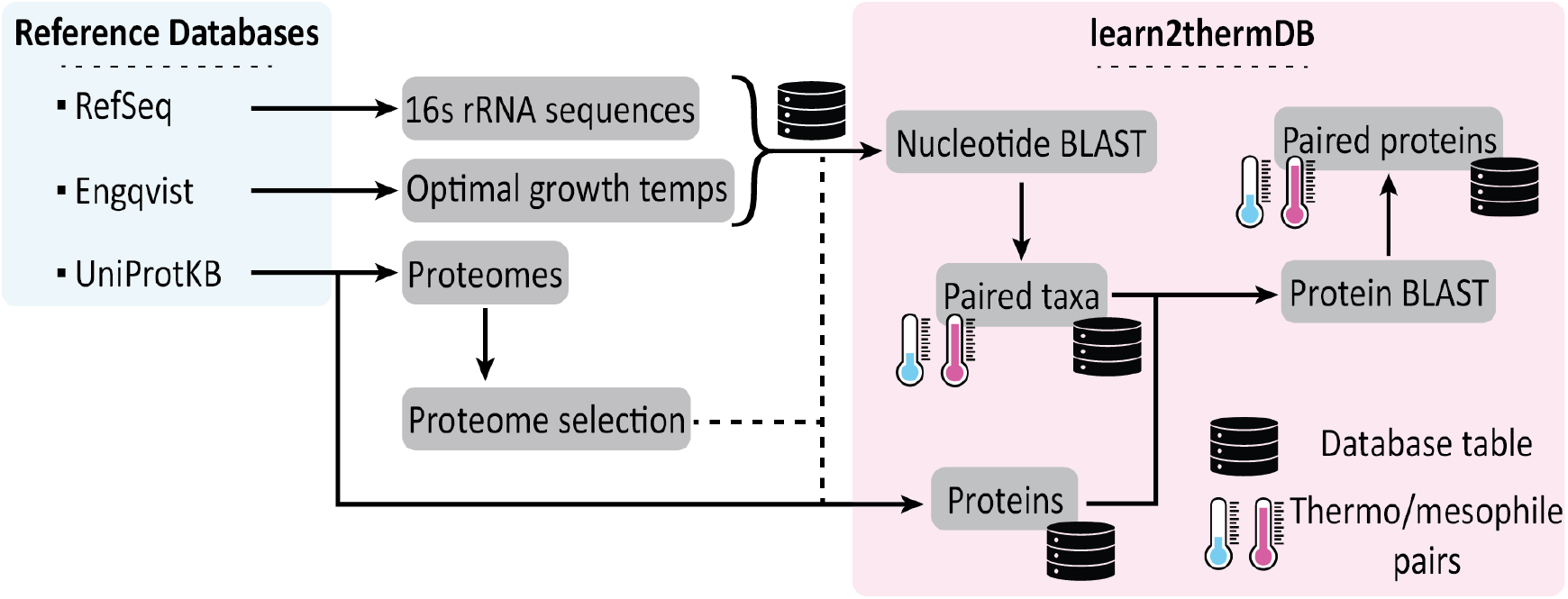
Workflow for labelling homologous protein pairs across temperature. Raw data includes RefSeq 16s rRNA sequences, OGT labels from Engqvist, UniProtKB proteome metadata and proteins. Proteome metadata is parsed to identify a single proteome for highly represented organisms, while retaining data for weakly studied taxa. Proteins are filtered such that only ones from the chosen proteomes and for which we have OGT are kept. Protein pair search space is filtered by first identifying pairs of related organisms via 16s rRNA alignment. Protein pairs are searched for by alignment of sequences. Final database tables are taxa, pairs of meso/thermo taxa, proteins, and pairs of meso/thermo proteins.

### Ingestion of raw data records

We retrieved Archaeal and Bacterial 16s rRNA sequences, along with their associated NCBI taxonomy identifier (taxid) from NCBI BioProjects 33175 and 33317 using the Entrez API.^66,67^ We only retained full sequences, ranging from 1300 to 1600 bases. When multiple sequences mapped to the same taxid, we observed >98% identity and retained the longest. Curated OGTs, produced by Engqvist, were then downloaded.^68,69^ For cases with multiple OGTs corresponding to the same species taxid, we used the average. The OGTs and 16s rRNA sequences were subsequently linked via taxid, generating a table with both quantities measured. For the next steps, we categorised organisms with OGT >40 °C as thermophiles. It should be noted that this information is tracked, enabling the users to filter data using a more stringent thermophilic label, such as 60 °C.^3^

We downloaded Archaeal and Bacterial protein sequences from UniProtKB.^70^ UniProtKB Proteome metadata was also retrieved to minimise redundancy while preserving taxonomic diversity. For strains with multiple proteomes, if any, we selected a single proteome based on UniProt’s priority labels: “Reference and representative proteome”, “Reference proteome”, “Representative proteome”. If these labels were absent, we discarded “Redundant proteome” and “Excluded proteome”, choosing the most populous remaining proteome. This process was repeated at the species level. Protein data files were parsed for primary sequence, host organism taxid, PDB, and AlphaFold database ID. Proteins not mapping to an organism in the taxa table were discarded. If a protein mapped to a proteome, it was retained if it belonged to the priority proteome identified for the organism, or if the priority proteome for the organism contained fewer than 2000 proteins. This avoided dataset saturation with model organism proteins while still including novel or infrequently studied proteins.

### Filtering protein homologous pair search space

To provide context-dependent information on protein thermal stability, pairs of proteins must be identified from a large pool of thermophilic and mesophilic ones. To do this we identified homologous sequences using a BLAST-like local alignment.^71^ The search space, considering only proteins <250 amino acids, was approximately 4.5 trillion pairs. A full pairwise BLAST-like search given the protein data space was projected to cost an unacceptable amount of carbon at 50,000 kg. To mitigate this, we first filtered the search space by identifying evolutionarily related taxa across temperatures. Using BLASTn, we aligned 16s rRNA sequences for mesophilic and thermophilic organisms. This sequence evolves slowly and is frequently used for inferring taxonomic relationships.^72–74^ See Supplementary Information S3 for alignment parameters. Any organism with >81% gap compressed sequence identity and >98.5% coverage on both strands was considered a taxa pair for the subsequent protein homolog searches, resulting in 150k taxa pairs and a search space of 230 billion possible protein pairs. The taxonomic breakdown and protein contributions of these pairs are depicted in Fig. 1-left.

### Identifying protein homologs across temperature

Using DIAMOND, we executed pairwise local sequence alignments for each thermophilic-mesophilic taxa pair, considering only proteins of 250 amino acids or less.^75^ A resource test indicated that DIAMOND had a twofold speed increase, while preserving sensitivity (see Supplementary Information S4 for alignment parameters and a comparison with BLASTp). Several alignment metrics such as percent identity and alignment coverage were monitored (refer to Supplementary Information S2 for a comprehensive list and definitions). This process yielded 69 million putative protein pairs with an E-value less than 1e-4 and over 75% coverage of both sequences. Using a stricter definition of thermophilicity (OGT >60 °C), the number of protein pairs reduces to 9 million. Despite this reduction, the dataset remains significantly larger than any existing homologous sequence collection across temperature.^48^ The protein space occupied by protein pairs is depicted in Figure 1-right, covering much of known protein space.

### Data pipeline

The entire process of developing the dataset, from extraction of raw data to downstream validation steps (see Technical Validation), is tracked using data version control (DVC).^76^ Thus, the complete history of how the data changed as the code was developed and the parameters were changed (eg. maximum protein length, minimum alignment metrics etc.) is available. Additionally, the pipeline can be rerun with a single command after environment and computing cluster configuration. The data tables created in this process (taxa, proteins, taxa_pairs, and pairs) are collected and linked as a relational database using DuckDB, allowing for fast access and filtering of the data.^77^ A depiction of the data pipeline is shown in Figure 3. A key subset of tunable parameters is shown in Table 3 along with the values used for the presented data. We hope that this organisation and data transparency will allow others to experiment with the data pipeline to suit their needs. The carbon cost of compute consuming steps in the pipeline was estimated using CodeCarbon.^65^ A comprehensive description of the pipeline steps and parameters is given in Supplementary Information S1.

**Table 3:**
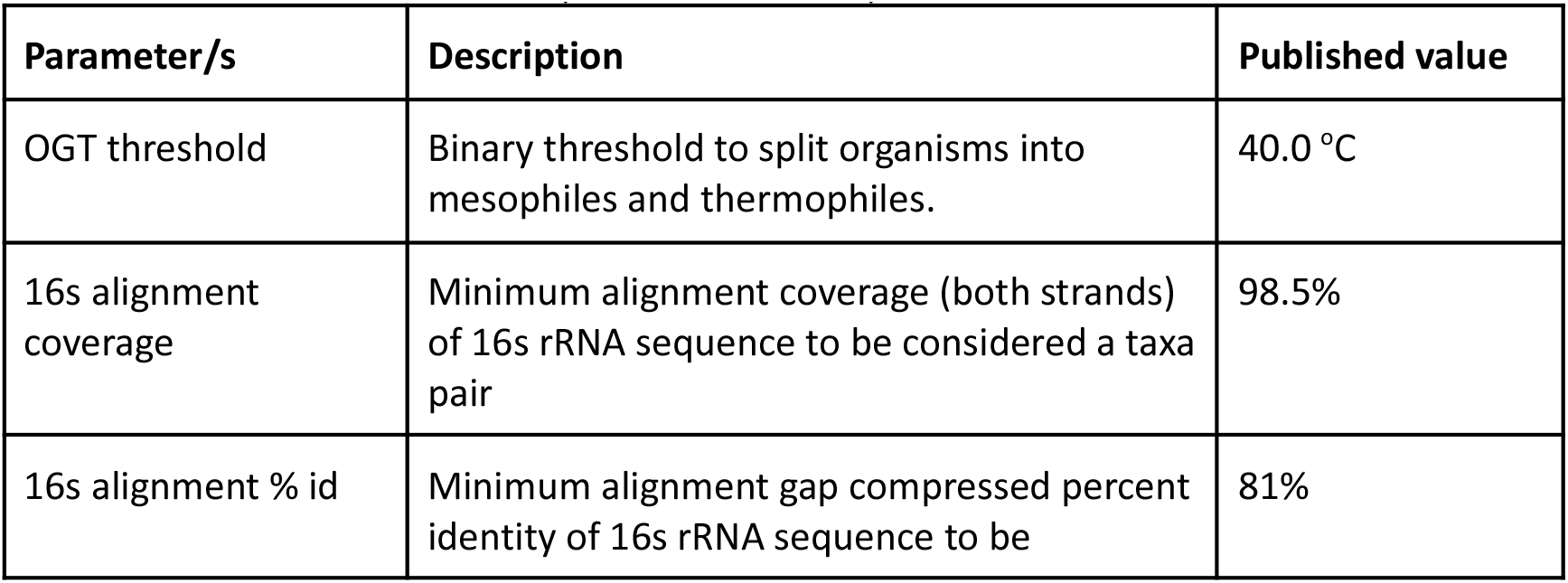
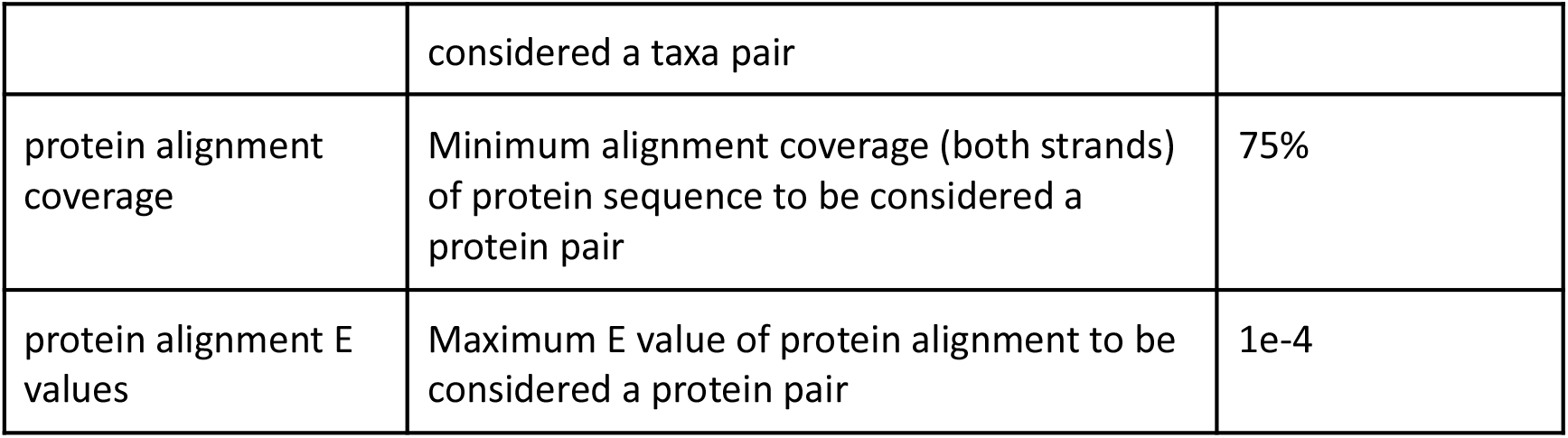
A small selection of tunable parameters used to produce the final dataset.

## Data Records

The dataset is available on Figshare^62^ in the form of a relational database, as well as a dump of semicolon separated value files.

### Schema

The dataset is structured as a relational database, implemented with DuckDB, an analytical database management system.^77^ It consists of four main tables: ‘taxa’, ‘proteins’, ‘taxa_pairs’ and (protein) ‘pairs’. An abbreviated schema detailing the relationships between these tables is illustrated in Figure 4. For a more detailed schema and a comprehensive description of each field, please refer to Supplementary Information S5.

**Figure 4.**
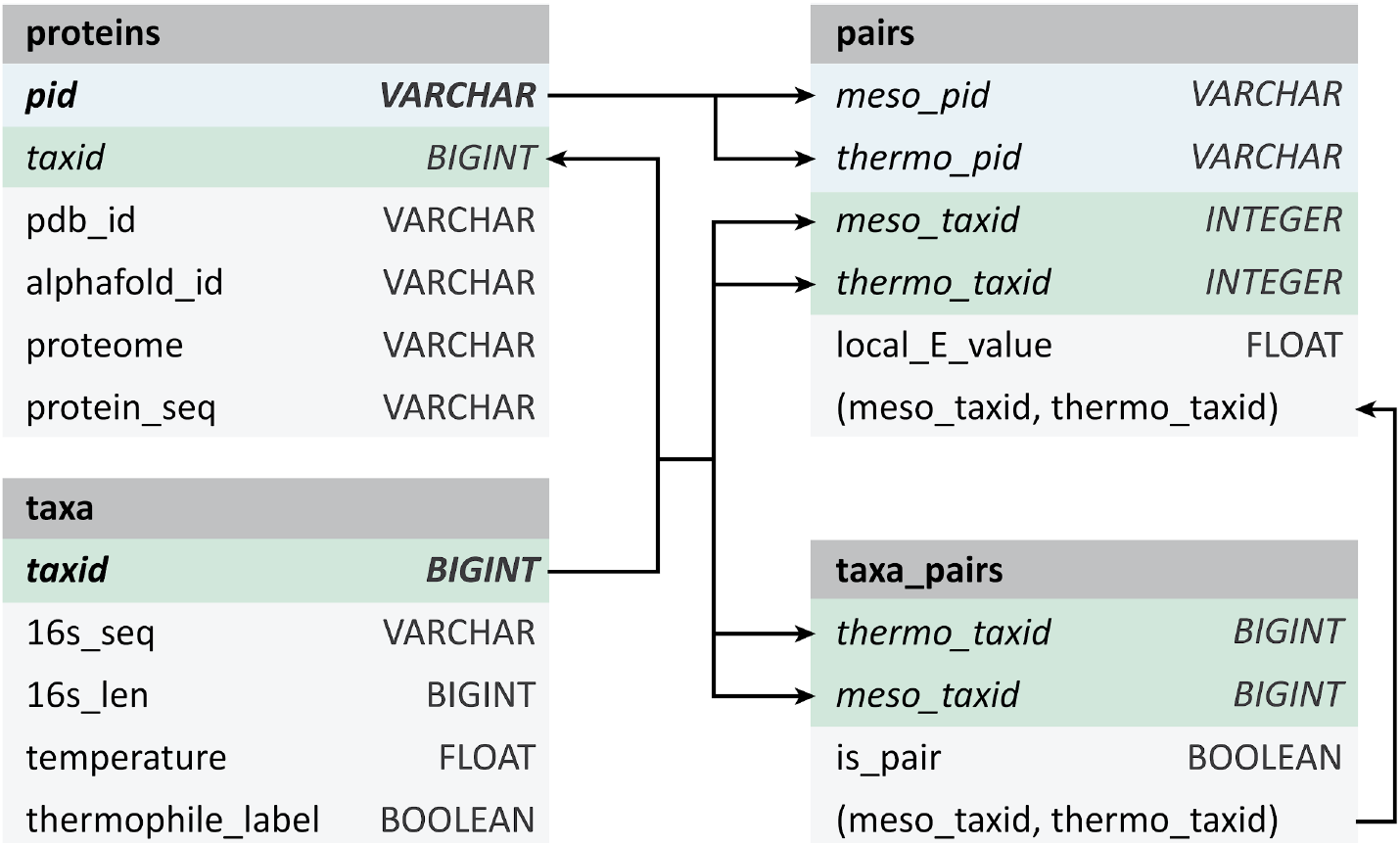
Abbreviated schema for the presented database. The taxa table contains NCBI taxonomic information, from superkingdom to species, as well as 16s rRNA sequence and OGT. The protein table contains protein sequence and external database links. The taxa_pairs and pairs table contain metrics for alignment for 16s rRNA sequences and protein sequences, respectively.

For 16s rRNA alignment (table “taxa_pairs”) and protein alignment (table “pairs”) conducted using DIAMOND, various metrics including bit score, coverage, etc. are reported and referenced throughout the manuscript. Detailed definitions of these metrics are provided in Supplementary Information S2.

### Accessing the dataset

To access the dataset, refer to the provided link on Figshare.^62^ The dataset can be found in a zipped file named “database.tar.gz.” This file includes the database “learn2therm.ddb”, a minimal environment file for querying the database called “enviroment.yml,” and a set of instructions provided in the “README.md” file. The database employs SQL, with some specific elements defined by DuckDB. For example, to retrieve all protein pairs and their corresponding amino acid sequences with >95% alignment coverage of both strands and thermophilic temperature >80 °C, use the following query:

SELECT m.protein_seq AS meso_seq, t.protein_seq AS thermo_seq

FROM pairs

INNER JOIN proteins AS m ON (m.pid=pairs.meso_pid)

INNER JOIN proteins AS t ON (t.pid=pairs.thermo_pid)

INNER JOIN taxa ON (taxa.taxid=pairs.thermo_taxid)

WHERE pairs.query_align_cov>0.95

AND pairs.subject_align_cov>0.95

AND taxa.temperature>80.0

This returns protein pairs in <10 seconds. See the README in the zipped data file for some additional example queries. Duckdb allows for exporting to csv, parquet, and many other desired formats, thus a user can retrieve the specific data they require on a taxa, protein, or protein pair basis.^77^

We have also provided dumps of the database tables as semicolon separated value files, however they may be unwieldy and it is recommended to use the database interface.

## Technical Validation

### Mapping to existing data

To ensure a proper mapping of OGT label to protein, we have joined our proteins with enzymes from Engqvist et al.^69^ via UniProt ID. Comparing the labels for the records in both datasets (N=1.6 mil) yields an R^2^ of 0.995. Some small differences in OGT labels arise due to our strain aggregation procedure, described above.

### Growth temperature as a proxy for melting temperature

To validate that the optimal growth temperature (OGT) is a suitable substitute for melting temperature as a measure of thermostability, and to ensure that temperature data has been accurately mapped, we compared proteins within our dataset to existing melting temperature datasets. We paired wild type proteins from FireProtDB and the Meltome atlas to proteins within our dataset using >99% coverage and identity, yielding 4,640 proteins with both internal OGT labels and external T_M_ labels.^43,45^ A Spearman’s correlation of 0.85 with a p-value of 0.0 was observed between these two quantities, suggesting a strong correlation. Furthermore, a binomial test was performed to determine if the melting temperature has a >99% chance of being greater than OGT, yielding a P-value of 2.68e-19. Figure 5 provides a parity plot of these two values. This analysis indicates that the trend observed in an organism’s protein melting temperatures is reflected in its growth temperature.

**Figure 5.**
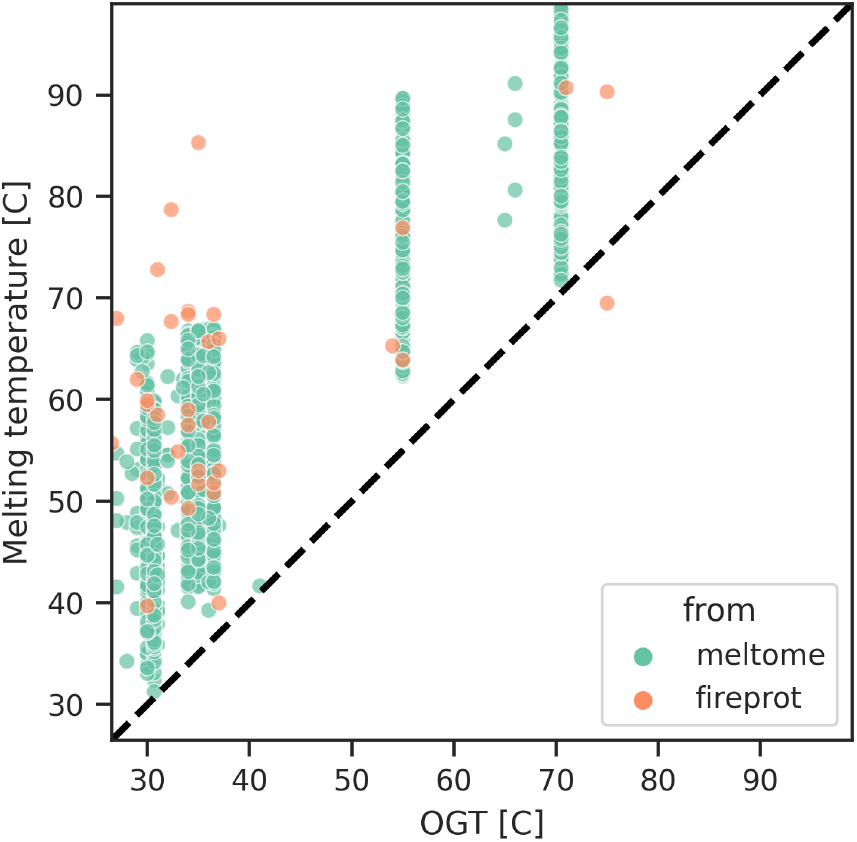
Melting temperature vs OGT, using data from two third party databases. The black dashed line is identity, and almost all examples fall on the side of T_M_ > OGT. The adage that melting temperature is greater than OGT is clearly supported, with a Spearman’s of 0.85 (P value 0.0), and a binomial test of >99% chance of passing with alternative P-value 2.68e^-19^.

### Comparing to alignments of known functional pairs

Hait et al. previously produced the current largest dataset of functional protein pairs across temperature at 1.6k pairs by starting with structures of PDB entries.^48^ This dataset served as our benchmark, with the quality of their protein pairs used for comparison. To facilitate this comparison, we first aligned their dataset using DIAMOND, and subsequently compared the resulting alignment metrics to ours.^75^ Statistically similar or even superior alignment scores to the baseline were observed in our dataset, as depicted in Table 4. Distributions of alignment percent identity and homology (as indicated by normalised bit score) are presented for both our data and the baseline pairs in Figure 6a-b. Further comparisons given in the table and figure are discussed in the subsequent sections.

**Table 4:** Statistical comparison of learn2therm protein pairs to Hait et al.’s pairs. The base set of protein pairs (alignment coverage >75%) as well as a subset (N=24 mil, coverage >95%) are shown. Percent identity, bit score (normalised to the average length of both proteins), and Pfam annotation Jaccard score are t-tests, while output P-value of structural alignment was treated as a binary if smaller than 0.001. For all metrics, our pairs meet the baseline on average for the subset with >90% alignment coverage, and most are still quality for the full set. Statistical significance is bolded.

| Score | Statistical Test | Probability (Cov>75) | Probability (Cov>95) |
| --- | --- | --- | --- |
| Normalised percent identity | left t | **$1.75e^{-6}$** | **$6.06e^{-155}$** |
| Normalised bit score | left t | **$1.94e^{-8}$** | **$9.47e^{-117}$** |
| Pfam Jaccard | left t | 0.99 | **$3.24e^{-13}$** |
| Structural P score | binomial<br>P<0.001 | **<0.001** | **<0.001** |

**Figure 6.**
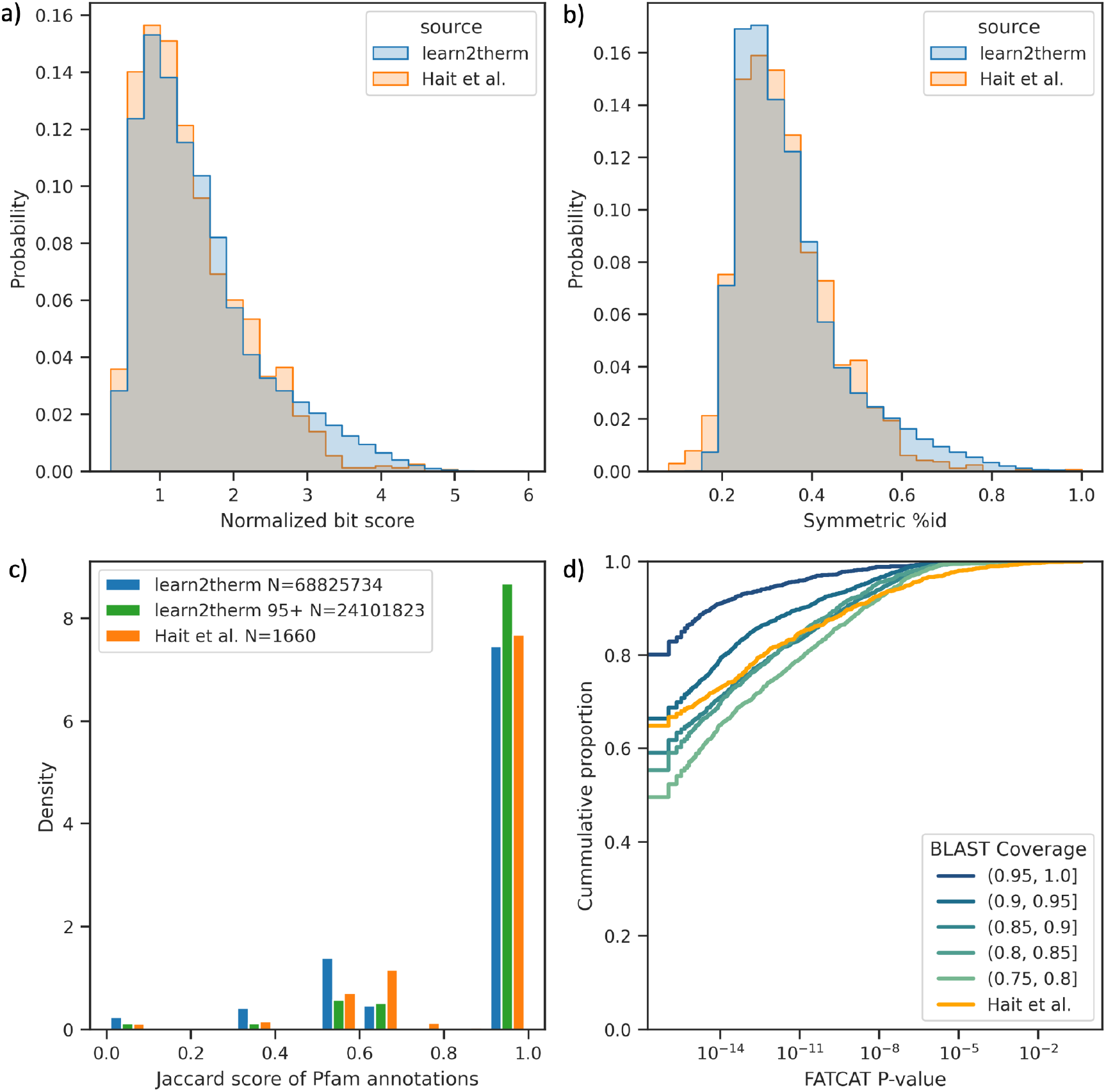
Comparison of protein pair quality between our data and Hait et al.’s 1660 protein pairs.^48^ A) Empirical distribution of local alignment homology as bit score normalised to the average length of both protein strands. Our data has a right shifted score on average with t-test probability=1.94e^-8^. B) Same as A, except with percent identity. Our data has a right shifted score on average with t-test probability=1.75e^-6^. C) Empirical distribution of Jaccard score over Pfam annotations for our data (blue) compared to the baseline data (orange). Our full data has more annotation mismatches on average. When only the 25 mil protein pairs with BLAST coverage >95% are considered, the Pfam annotations become indistinguishable from the baseline with t-test probability=3.24e^-13^. D) Cumulative distribution of FATCAT structural alignment P-value for bins in BLAST coverage uniformly sampled from our data, compared to the baseline structural alignments. Even low coverage pairs are more likely to have less than one in a thousand P-value with binomial confidence >99%.

### Pfam annotations

We used Pfam to annotate proteins in both our dataset and Hait’s pairs.^48,78–80^ Retaining matches with an E-value < 1e-10 (normalised using the size of Pfam 35.0) resulted in 86.1% of our proteins and 99.8% of Hait’s proteins being labelled with at least one annotation. This suggests that our data includes novel proteins not extensively represented in Pfam. We evaluated the quality of a protein pair according to Pfam by calculating the set Jaccard score of accession annotations (Supplementary Information Eq. 1). We only considered pairs where at least one member was labelled with at least one annotation, as the remaining are out of scope for Pfam. Although our data contains slightly more annotation mismatches, if only the N=24 million pairs with >95% sequence alignment coverage are considered, the scores are indistinguishable from the baseline pairs, with a t-test probability of 3.24e^-13^. The score distributions for both datasets are compared in Figure 6-C. A detailed description of this search is available in Supplementary Information S6. Homology searches using HMMs are highly sensitive to evolutionary distance, suggesting that true pairs of proteins with the same or similar functions are likely to share Pfam annotations, if available.^81^

### Structural alignments

We used FATCAT to align PDB structure of the baseline pairs with flexible alignment.^82,83^ Chain A was chosen if present. This flexible protein structure alignment algorithm provides an empirically scaled score “P-value” which indicates the likelihood that the raw alignment score were to occur between two random proteins, thus a P-value close to 0.0 indicates a quality structural overlap. We repeated this process for our protein pairs using PDB structures where available, otherwise utilising AlphaFold predicted structures.^57,84^ For learn2therm protein pairs, we took a subset of 10k pairs due to computational cost limitations of structural alignment. We took this sample uniformly across 5 bins in sequence alignment coverage between 75% (the dataset minimum) and 100%. The cumulative distribution of probability scores from FATCAT for each of these subsets is given in Figure 6-D, where we see that our protein pairs have alignments even less likely to be observed randomly than the already quality baseline. To compare the alignment results to the baseline statistically, we considered a binary problem where FATCAT probability of occurring randomly less than one in a thousand is considered a quality protein pair. Conducting a binomial test, even pairs with sequence alignment coverage <80% are indistinguishable from the baseline, and higher alignment coverage yields structural alignment with smaller P-values than Hait’s on average with >99.9% confidence.

### Signal of growth temperature predictors

In order to evaluate the signal for downstream deep learning models on our data, we consider a classifier of thermophilic versus mesophilic origin from protein sequence alone. This is one of the simplest models that can be produced from our data, and other work has shown this to be a learnable function.^46,47,85–87^ To facilitate this test, we preprocess our data using the following steps: binarization by OGT <30 °C or >=60 °C, class balancing, deduplication of similar sequences, and splitting based on NCBI taxonomy of host species, resulting in a training and test set of 290k and 28k proteins respectively, each labelled as mesophilic or thermophilic (see Supplementary Information S7 for details on the rigorous preprocessing conducted).

We then evaluated a recent predictor, TemStaPro, on our test set of 28k proteins.^47^ This model is an ensemble of neural networks trained on top of the output embeddings of ProtT5XL, a Large Protein Language Model (LPLM).^88^ The TemStaPro ensemble outputs predictions in bins of 5 °C between ‘<40’ and ‘65<=‘, and conducts a self consistency check between ensembles. On our data, the model is consistent for 95.5% of examples, and of those, predicts the correct class with 91% accuracy compared to the null model of predicting the majority class with 61% accuracy. The distribution of predictions is shown in Figure 7 below, where the model is clearly a predictor of >=60 and <30 °C labels.

**Figure 7.**
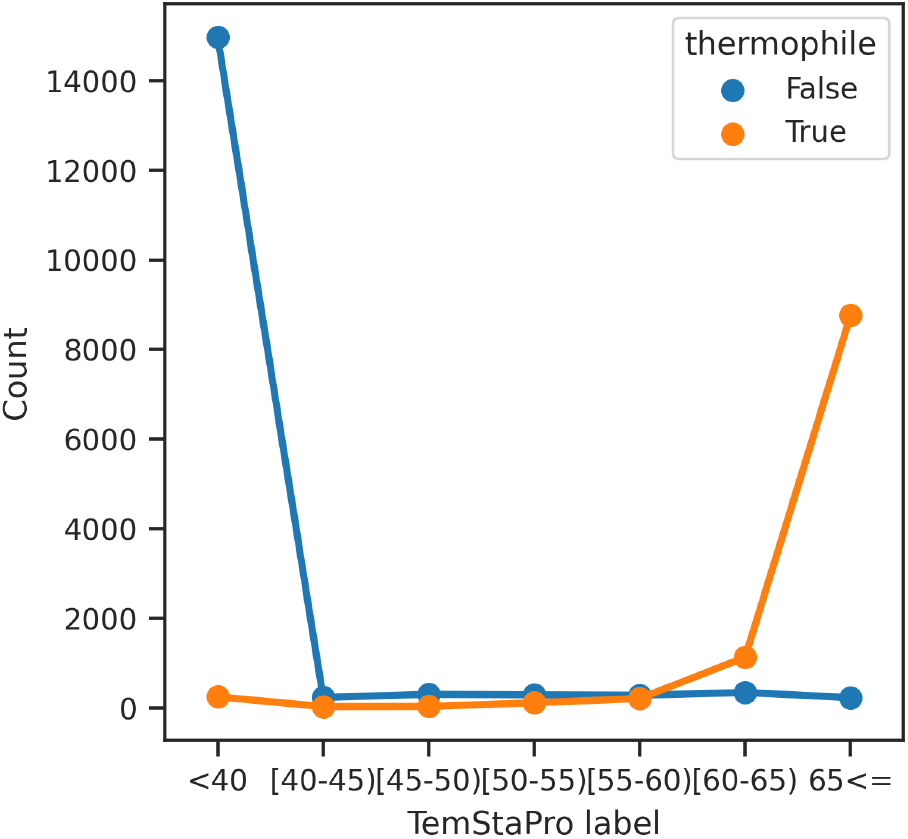
Predictions on our test set by TemStaPro.^47^ Thermophilic label is >=60 °C versus <30 °C. The model makes very few predictions between these two quantities where there is no data, as expected, and is able to recover the actual classes with 91% accuracy.

Given that TemStaPro was trained on UniParc proteins, it is possible that the model has seen data from our test set before. To ensure that high performance is not due to data leakage, we trained our own model on the development set of our data. We finetuned ProteinBERT, a LPLM, on our development set.^88^ The training parameters and architectural details can be found in Supplementary Information S7. The performance on our held out test set is shown in Table 5 below. The model is highly predictive, confirming that our dataset of proteins and OGTs retain signal.

**Table 5:** Test set performance of our fine tuned LPLM.

| Balanced Accuracy | F1 Score | Matthew's Correlation | Confusion Matrix |
| --- | --- | --- | --- |
| 92.5% | 0.91 | 0.85 | [16069, 1459, 671, 10296] |

It is clear that the data we produced is capable of supporting the foundational machine learning task of thermophilic classification, yet with almost an order of magnitude more data than previously. With the addition of homologous pairing, as well as its increased size, the dataset will open the door for more complex models such as thermal stability design models.

## Usage Notes

The codebases that produced the results in this work are free and openly available. The pipeline leverages Data Version Control (DVC).^76^ A user can reproduce this data with a single command ‘dvc exp run’ (assuming available compute resources) or run the data pipeline using different parameters and produce a variant of the dataset, all the while monitoring how the data changes. Environments and parameters necessary for this process are retained within the repositories.

A snapshot of the data repositories at the time of production of these results is provided as a Figshare dump in addition to the repository links. Each of these repositories is DVC tracked, thus the set of parameters used to execute the pipeline is found in ‘params.yaml’. A description of each parameter is given as comments in the file, and in Supplementary Information S1. See Table 6 for the repositories and data dumps. In order to easily produce a directed acyclic graph of steps within the pipelines, dependencies and outputs of each pipeline stage, and view experiment results as parameters were changed, see DVC’s API.^76^

**Table 6:**
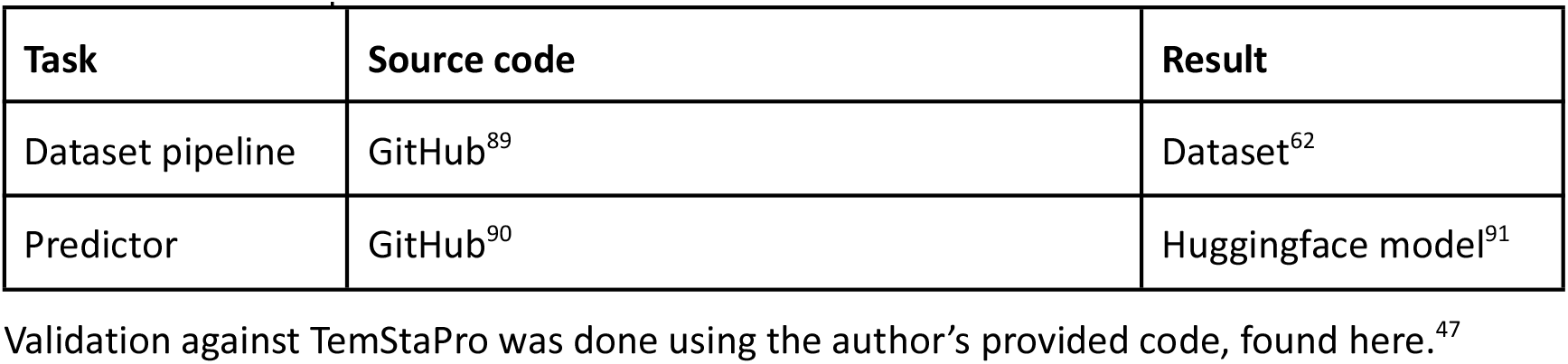
Location of open code and data.

## Code Availability

All of the products of this manuscript as well as the parameterized code pipelines are free and openly available. Please see “Usage Notes.”

## Acknowledgements

This work was funded by NSF Engineering Data Science Institute Grant OAC-1934292.

The research team would like to acknowledge the computer resources and IT team at UW’s Hyak supercomputer.

The University of Washington acknowledges the Coast Salish peoples of the land where this work was conducted, the land which touches the shared waters of all tribes and bands within the Suquamish, Tulalip and Muckleshoot nations.

## Author contributions

EK designed the pipeline for the task of producing protein pairs, wrote the code for the pipeline, conducted analysis, and wrote the manuscript.

HA wrote code components for scanning the database with pyhmmer, and wrote the manuscript.

RF analysed data and created figures within the manuscript. CV wrote code components for running FATCAT on our data.

DACB designed the task for finding protein pairs, and wrote the manuscript.

EK, HA, RF, CV, LR, AM, and DACB contributed to brainstorming and editing the manuscript.

## Competing interests

The authors claim no competing interests.

## Notes

https://doi.org/10.6084/m9.figshare.23589390.v1

https://doi.org/10.6084/m9.figshare.23581932.v1

https://doi.org/10.6084/m9.figshare.23589210.v1

https://doi.org/10.6084/m9.figshare.23582325.v1

### Competing Interest Statement

The authors have declared no competing interest.

## References

1. Narasimhan, D. et al. Structural analysis of thermostabilizing mutations of cocaine esterase. Protein Eng. Des. Sel. 23, 537–547 (2010).

2. Xiong, X. et al. A thermostable, closed SARS-CoV-2 spike protein trimer. Nat. Struct. Mol. Biol. 27, 934–941 (2020).

3. Mehta, R., Singhal, P., Singh, H., Damle, D. & Sharma, A. K. Insight into thermophiles and their wide-spectrum applications. 3 Biotech 6, 81 (2016).

4. Kumar, V., Marín-Navarro, J. & Shukla, P. Thermostable microbial xylanases for pulp and paper industries: trends, applications and further perspectives. World J. Microbiol. Biotechnol. 32, 34 (2016).

5. Knott, B. C. et al. Characterization and engineering of a two-enzyme system for plastics depolymerization. Proc. Natl. Acad. Sci. 117, 25476–25485 (2020).

6. Polizzi, K. M., Bommarius, A. S., Broering, J. M. & Chaparro-Riggers, J. F. Stability of biocatalysts. Curr. Opin. Chem. Biol. 11, 220–225 (2007).

7. Berezovsky, I. N., Zeldovich, K. B. & Shakhnovich, E. I. Positive and Negative Design in Stability and Thermal Adaptation of Natural Proteins. PLOS Comput. Biol. 3, e52 (2007).

8. Modarres, H. P., Mofrad, M. R. & Sanati-Nezhad, A. Protein thermostability engineering. RSC Adv. 6, 115252–115270 (2016).

9. Åqvist, J., Isaksen, G. V. & Brandsdal, B. O. Computation of enzyme cold adaptation. Nat. Rev. Chem. 1, 1–14 (2017).

10. Tokuriki, N. & Tawfik, D. S. Stability effects of mutations and protein evolvability. Curr. Opin. Struct. Biol. 19, 596–604 (2009).

11. Atsavapranee, B., Stark, C. D., Sunden, F., Thompson, S. & Fordyce, P. M. Fundamentals to function: Quantitative and scalable approaches for measuring protein stability. Cell Syst. 12, 547–560 (2021).

12. Berezovsky, I. N. & Shakhnovich, E. I. Physics and evolution of thermophilic adaptation. Proc. Natl. Acad. Sci. 102, 12742–12747 (2005).

13. Takano, K., Aoi, A., Koga, Y. & Kanaya, S. Evolvability of Thermophilic Proteins from Archaea and Bacteria. Biochemistry 52, 4774–4780 (2013).

14. Sawle, L. & Ghosh, K. How Do Thermophilic Proteins and Proteomes Withstand High Temperature? Biophys. J. 101, 217–227 (2011).

15. England, J. L., Shakhnovich, B. E. & Shakhnovich, E. I. Natural selection of more designable folds: A mechanism for thermophilic adaptation. Proc. Natl. Acad. Sci. 100, 8727–8731 (2003).

16. Traxlmayr, M. W. & Shusta, E. V. Directed Evolution of Protein Thermal Stability Using Yeast Surface Display. in Synthetic Antibodies: Methods and Protocols (ed. Tiller, T.) 45–65 (Springer, 2017).

17. Akram, F., Haq, I. ul, Aqeel, A., Ahmed, Z. & Shah, F. I. Thermostable cellulases: Structure, catalytic mechanisms, directed evolution and industrial implementations. Renew. Sustain. Energy Rev. 151, 111597 (2021).

18. Zhao, H. & Arnold, F. H. Directed evolution converts subtilisin E into a functional equivalent of thermitase. Protein Eng. Des. Sel. 12, 47–53 (1999).

19. Huang, J. X. et al. High throughput discovery of functional protein modifications by Hotspot Thermal Profiling. Nat. Methods 16, 894–901 (2019).

20. Pongpamorn, P. et al. Identification of a Hotspot Residue for Improving the Thermostability of a Flavin-Dependent Monooxygenase. ChemBioChem 20, 3020–3031 (2019).

21. Son, H. F. et al. Rational Protein Engineering of Thermo-Stable PETase from Ideonella sakaiensis for Highly Efficient PET Degradation. ACS Catal. 9, 3519–3526 (2019).

22. Merkley, E. D., Parson, W. W. & Daggett, V. Temperature dependence of the flexibility of thermophilic and mesophilic flavoenzymes of the nitroreductase fold. Protein Eng. Des. Sel. 23, 327–336 (2010).

23. Pikkemaat, M. G., Linssen, A. B. M., Berendsen, H. J. C. & Janssen, D. B. Molecular dynamics simulations as a tool for improving protein stability. Protein Eng. Des. Sel. 15, 185–192 (2002).

24. Packer, M. S. & Liu, D. R. Methods for the directed evolution of proteins. Nat. Rev. Genet. 16, 379–394 (2015).

25. Defresne, M., Barbe, S. & Schiex, T. Protein Design with Deep Learning. Int. J. Mol. Sci. 22, 11741 (2021).

26. Wang, J., Cao, H., Zhang, J. Z. H. & Qi, Y. Computational Protein Design with Deep Learning Neural Networks. Sci. Rep. 8, 6349 (2018).

27. Linder, J., Bogard, N., Rosenberg, A. B. & Seelig, G. A Generative Neural Network for Maximizing Fitness and Diversity of Synthetic DNA and Protein Sequences. Cell Syst. 11, 49–62.e16 (2020).

28. Ding, W., Nakai, K. & Gong, H. Protein design via deep learning. Brief. Bioinform. 23, bbac102 (2022).

29. Watson, J. L. et al. De novo design of protein structure and function with RFdiffusion. Nature (2023) doi:10.1038/s41586-023-06415-8.

30. Dauparas, J. et al. Robust deep learning based protein sequence design using ProteinMPNN. 2022.06.03.494563 Preprint at 10.1101/2022.06.03.494563 (2022).

31. Syrlybaeva, R. & Strauch, E.-M. Deep learning of protein sequence design of protein–protein interactions. Bioinformatics 39, btac733 (2023).

32. Kuhlman, B. Designing protein structures and complexes with the molecular modeling program Rosetta. J. Biol. Chem. 294, 19436–19443 (2019).

33. Kaufmann, K. W., Lemmon, G. H., DeLuca, S. L., Sheehan, J. H. & Meiler, J. Practically Useful: What the Rosetta Protein Modeling Suite Can Do for You. Biochemistry 49, 2987–2998 (2010).

34. Leman, J. K. et al. Macromolecular modeling and design in Rosetta: recent methods and frameworks. Nat. Methods 17, 665–680 (2020).

35. PDB Statistics: PDB Data Distribution by Natural Source Organism. RCSB Protein Data Bank https://www.rcsb.org/stats/distribution-source-organism-natural.

36. Casadio, R., Savojardo, C., Fariselli, P., Capriotti, E. & Martelli, P. L. Turning Failures into Applications: The Problem of Protein ΔΔG Prediction. in Data Mining Techniques for the Life Sciences (eds. Carugo, O. & Eisenhaber, F.) 169–185 (Springer US, 2022).

37. Louis, B. B. V. & Abriata, L. A. Reviewing Challenges of Predicting Protein Melting Temperature Change Upon Mutation Through the Full Analysis of a Highly Detailed Dataset with High-Resolution Structures. Mol. Biotechnol. 63, 863–884 (2021).

38. Nguyen, V. et al. Evolutionary drivers of thermoadaptation in enzyme catalysis. Science 355, 289–294 (2017).

39. Leuenberger, P. et al. Cell-wide analysis of protein thermal unfolding reveals determinants of thermostability. Science 355, eaai7825 (2017).

40. Ponnuswamy, P. K., Muthusamy, R. & Manavalan, P. Amino acid composition and thermal stability of proteins. Int. J. Biol. Macromol. 4, 186–190 (1982).

41. Karshikoff, A., Nilsson, L. & Ladenstein, R. Rigidity versus flexibility: the dilemma of understanding protein thermal stability. FEBS J. 282, 3899–3917 (2015).

42. Quezada, A. G. et al. Interplay between Protein Thermal Flexibility and Kinetic Stability. Structure 25, 167–179 (2017).

43. Jarzab, A. et al. Meltome atlas-thermal proteome stability across the tree of life. Nat. Methods 17, 495–503 (2020).

44. Nikam, R., Kulandaisamy, A., Harini, K., Sharma, D. & Gromiha, M. M. ProThermDB: thermodynamic database for proteins and mutants revisited after 15 years. Nucleic Acids Res. 49, D420–D424 (2021).

45. Stourac, J. et al. FireProtDB: database of manually curated protein stability data. Nucleic Acids Res. 49, D319–D324 (2021).

46. Li, G., Rabe, K. S., Nielsen, J. & Engqvist, M. K. M. Machine Learning Applied to Predicting Microorganism Growth Temperatures and Enzyme Catalytic Optima. ACS Synth. Biol. 8, 1411–1420 (2019).

47. Pudžiuvelytė, I. et al. TemStaPro: protein thermostability prediction using sequence representations from protein language models. 2023.03.27.534365 Preprint at 10.1101/2023.03.27.534365 (2023).

48. Hait, S., Mallik, S., Basu, S. & Kundu, S. Finding the generalized molecular principles of protein thermal stability. Proteins Struct. Funct. Bioinforma. 88, 788–808 (2020).

49. Pucci, F., Schwersensky, M. & Rooman, M. Artificial intelligence challenges for predicting the impact of mutations on protein stability. Curr. Opin. Struct. Biol. 72, 161–168 (2022).

50. Gromiha, M. M., Oobatake, M. & Sarai, A. Important amino acid properties for enhanced thermostability from mesophilic to thermophilic proteins. Biophys. Chem. 82, 51–67 (1999).

51. Miotto, M. et al. Insights on protein thermal stability: a graph representation of molecular interactions. Bioinformatics 35, 2569–2577 (2019).

52. Dehouck, Y., Folch, B. & Rooman, M. Revisiting the correlation between proteins’ thermoresistance and organisms’ thermophilicity. Protein Eng. Des. Sel. 21, 275–278 (2008).

53. Ahmed, Z., Zulfiqar, H., Tang, L. & Lin, H. A Statistical Analysis of the Sequence and Structure of Thermophilic and Non-Thermophilic Proteins. Int. J. Mol. Sci. 23, 10116 (2022).

54. Pucci, F. & Rooman, M. Improved insights into protein thermal stability: from the molecular to the structurome scale. Philos. Trans. R. Soc. Math. Phys. Eng. Sci. 374, 20160141 (2016).

55. Pucci, F. & Rooman, M. Physical and molecular bases of protein thermal stability and cold adaptation. Curr. Opin. Struct. Biol. 42, 117–128 (2017).

56. Jung, F., Frey, K., Zimmer, D. & Mühlhaus, T. DeepSTABp: A Deep Learning Approach for the Prediction of Thermal Protein Stability. Int. J. Mol. Sci. 24, 7444 (2023).

57. Jumper, J. et al. Highly accurate protein structure prediction with AlphaFold. Nature 596, 583–589 (2021).

58. Nambiar, A. et al. Transforming the Language of Life: Transformer Neural Networks for Protein Prediction Tasks. http://biorxiv.org/lookup/doi/10.1101/2020.06.15.153643 (2020) xdoi:10.1101/2020.06.15.153643.

59. Verkuil, R. et al. Language models generalize beyond natural proteins. 2022.12.21.521521 Preprint at 10.1101/2022.12.21.521521 (2022).

60. Anishchenko, I. et al. De novo protein design by deep network hallucination. Nature 600, 547–552 (2021).

61. AlQuraishi, M. & Sorger, P. K. Differentiable biology: using deep learning for biophysics-based and data-driven modeling of molecular mechanisms. Nat. Methods 18, 1169–1180 (2021).

62. Komp, E. et al. learn2thermDB. (Figshare, 2023). doi:10.6084/m9.figshare.23581932.

63. NCBI Taxonomy: a comprehensive update on curation, resources and tools | Database | Oxford Academic. https://academic.oup.com/database/article/doi/10.1093/database/baaa062/5881509?login=false.

64. Lin, Z. et al. Evolutionary-scale prediction of atomic-level protein structure with a language model. Science 379, 1123–1130 (2023).

65. CodeCarbon. Estimation of Computation Carbon Cost https://codecarbon.io/.

66. O’Leary, N. A. et al. Reference sequence (RefSeq) database at NCBI: current status, taxonomic expansion, and functional annotation. Nucleic Acids Res. 44, D733–D745 (2016).

67. Kans, J. Entrez Direct: E-utilities on the Unix Command Line. in Entrez Programming Utilities Help [Internet] (National Center for Biotechnology Information (US), 2023).

68. Engqvist, M. K. M. Growth temperatures for 21,498 microorganisms. (Zenodo, 2018). doi:10.5281/zenodo.1175609.

69. Engqvist, M. K. M. Correlating enzyme annotations with a large set of microbial growth temperatures reveals metabolic adaptations to growth at diverse temperatures. BMC Microbiol. 18, 177 (2018).

70. The UniProt Consortium. UniProt: a hub for protein information. Nucleic Acids Res. 43, D204–D212 (2015).

71. Camacho, C. et al. BLAST+: architecture and applications. BMC Bioinformatics 10, 421 (2009).

72. Yang, B., Wang, Y. & Qian, P.-Y. Sensitivity and correlation of hypervariable regions in 16S rRNA genes in phylogenetic analysis. BMC Bioinformatics 17, 135 (2016).

73. Schloss, P. D. The effects of alignment quality, distance calculation method, sequence filtering, and region on the analysis of 16S rRNA gene-based studies. PLoS Comput. Biol. 6, e1000844 (2010).

74. Kim, M., Oh, H.-S., Park, S.-C. & Chun, J. Towards a taxonomic coherence between average nucleotide identity and 16S rRNA gene sequence similarity for species demarcation of prokaryotes. Int. J. Syst. Evol. Microbiol. 64, 346–351 (2014).

75. Buchfink, B., Xie, C. & Huson, D. H. Fast and sensitive protein alignment using DIAMOND. Nat. Methods 12, 59–60 (2015).

76. Data Version Control · DVC. Data Version Control · DVC https://dvc.org/.

77. DuckDB | Proceedings of the 2019 International Conference on Management of Data. https://dl.acm.org/doi/abs/10.1145/3299869.3320212.

78. Mistry, J. et al. Pfam: The protein families database in 2021. Nucleic Acids Res. 49, D412–D419 (2021).

79. Finn, R. D., Clements, J. & Eddy, S. R. HMMER web server: interactive sequence similarity searching. Nucleic Acids Res. 39, W29–W37 (2011).

80. PyHMMER: a Python library binding to HMMER for efficient sequence analysis | Bioinformatics | Oxford Academic. https://academic.oup.com/bioinformatics/article/39/5/btad214/7131068.

81. Pearson, W. R. An Introduction to Sequence Similarity (“Homology”) Searching. Curr. Protoc. Bioinforma. Ed. Board Andreas Baxevanis Al 03, 10.1002/0471250953.bi0301s42 (2013).

82. Li, Z., Jaroszewski, L., Iyer, M., Sedova, M. & Godzik, A. FATCAT 2.0: towards a better understanding of the structural diversity of proteins. Nucleic Acids Res. 48, W60–W64 (2020).

83. Burley, S. K. et al./person-group>. Protein Data Bank (PDB): The Single Global Macromolecular Structure Archive. in Protein Crystallography: Methods and Protocols (eds. Wlodawer, A., Dauter, Z. & Jaskolski, M.) 627–641 (Springer, 2017). doi:10.1007/978-1-4939-7000-1_26.

84. AlphaFold Protein Structure Database: massively expanding the structural coverage of protein-sequence space with high-accuracy models | Nucleic Acids Research | Oxford Academic. https://academic.oup.com/nar/article/50/D1/D439/6430488.

85. Yang, Y., Zhao, J., Zeng, L. & Vihinen, M. ProTstab2 for Prediction of Protein Thermal Stabilities. Int. J. Mol. Sci. 23, 10798 (2022).

86. Wang, X.-F., Gao, P., Liu, Y.-F., Li, H.-F. & Lu, F. Predicting Thermophilic Proteins by Machine Learning. Curr. Bioinforma. 15, 493–502 (2020).

87. Zhao, J., Yan, W. & Yang, Y. DeepTP: A Deep Learning Model for Thermophilic Protein Prediction. Int. J. Mol. Sci. 24, 2217 (2023).

88. Elnaggar, A. et al. ProtTrans: Toward Understanding the Language of Life Through Self-Supervised Learning. IEEE Trans. Pattern Anal. Mach. Intell. 44, 7112–7127 (2022).

89. Komp, E., Alanzi, H., Vuong, C., Beck, D. & Francis, R. learn2thermDB data pipeline. (Figshare, 2023). doi:10.6084/m9.figshare.23589390.

90. Komp, E. & Beck, D. learn2thermML source code. (Figshare, 2023). doi:10.6084/m9.figshare.23589210.

91. Komp, Evan & Beck, David. learn2therm_model. (Huggingface, 2023). doi:10.57967/hf/0815.

