## Supplemental Information for "Homologous Pairs of Low and High Temperature Originating Proteins Spanning the Known Prokaryotic Universe"

### *S1. Pipeline steps and parameters*

The pipeline for downloading raw data, to validation of protein pairs is runnable with a single command: ‘dvc repro’. Alternatively you can run individual stages of the pipeline with ‘dvc repro -s <name_of_stage>’. Many of the stages in the pipeline have parameters that affect the runtime behaviour and stage outputs, thus the pipeline is tunable by modifying ‘params.yaml’. Each stage in the pipeline is outlined below, and the parameters are given in Table S1.

#### Data ingestion

1. **`s0.0_get_raw_data_taxa.py`**
   Pull most recent NCBI 16s r RNA sequences, and OGT records from Enqvist
   - Params: `min_16s_len`, `max_16s_len` number of nucleotides required to keep and organism
   - Outputs: `data/taxa.parquet`, columns include OGT, 16s sequence, taxid, other taxonomy
   - Metrics: `n_taxa` total number of labelled organisms, `taxa_pulled_date` when the data was retrieved
2. **`s0.1_get_raw_data_proteins.py`**
   Retrieve single cell uniprot. Uses FTP to download very large uniprot files
   - Inputs: `data/taxa.parquet`
   - Outputs: `data/taxa/uniprot/uniprot_pulled_timestamp` indicates when files were pulled
   - Metrics: `taxa_pulled_date` when data was retrieved
   - Untracked Outputs: The script produces `*.xml.gz` files that are untracked because they take so long to download. DVC ignores them, subsequent calls to the script skip downloading files already present.

3. **`s0.2_get_proteome_mdata.py`**
   Get metadata for UniProt proteomes. Selects one "best" proteome per organism.
   - Outputs: `data/uniprot/proteome_metadata.csv`, columns include taxa pair file, number of hits, emissions, total searchable space
4. **`s0.3_parse_proteins.py`**
   Extract minimal protein data and store in an efficient file format. Skip proteins that we don't have OGT for or are from redundant proteomes
   - Params: `max_prot_per_file` size of parquet files
   - Inputs: `data/taxa/uniprot/uniprot_pulled_timestamp`, `data/taxa.parquet`
   - Outputs: `data/proteins`, contains proteins in chunked files of the form `*.parquet`. Columns include protein sequence, database identifiers, and associated taxa IDs, `./data/metrics/s0.3_protein_per_data_distr.csv` table of number of proteins per taxa
   - Metrics: `n_proteins` total protein count, `percent_prot_w_struc` fraction of proteins with PDB or AlphaFold id

#### Data pairing

5. **`s1.0_label_taxa.py`**
   Assign booleans for taxa as thermophile
   - Params: `ogt_threshold` binary thermophile threshold
   - Inputs: `data/taxa.parquet`
   - Outputs: `data/taxa_thermophile_labels.parquet`
   - Metrics: `n_meso`, `n_thermo`
6. **`s1.1_get_16s_blast_scores.py`**
   Compute pairwise BLAST pairings of meso vs thermo 16s rRNA sequences.
   - Params: `16s_blast_parameters` (there are a number), `blast_metrics`
   - Inputs: `data/taxa*.parquet`, `./data/metrics/s0.3_protein_per_data_distr.csv` (used to skip alignment if taxa has no proteins)
   - Outputs: `data/taxa_pairs/alignment/*.parquet` table of taxids and BLAST scores.
7. **`s1.2_label_all_pairs.py`**
   Create a list of taxa pairs that meet a minimum 16s rRNA BLAST score.
   - Params: `blast_metric_thresholds` defines thresholds on 16s blast metrics to consider a pair
   - Inputs: `data/taxa_pairs/alignment/*.parquet`
   - Outputs: `data/taxa_pairs/pair_labels/*.parquet` 1:1 index mapping to data/taxa_pairs/alignment/*.parquet of boolean labels of whether that taxa are a pair
   - Metrics: `num_taxa_pairs_conservative` number of pairs that passed thresholds. Only this number will we blastp, `taxa_pair_found_ratio` fraction of pairs with metrics eg n_taxa_pairs/(n_therm*n_meso) that will be searched for protein pairs
8. **`s1.3_protein_alignment.py`**
   Runs a massive parallel cluster to align protein pairs among taxa pairs using DIAMOND.
   - Params: `dask_cluster_class` class for cluster in dask_jobqueue, `max_protein_length`, `method` local aligner type, `n_jobs` parallel workers, each doing a taxa pair at a time, `method_X_params` where X is eg. "blast" params given to aligner, `blast_metrics` alignment metrics to record.
   - Inputs: `data/taxa_pairs/alignment/*.parquet`, `data/taxa_pairs/pair_labels/*.parquet`, `./data/proteins`
   - Outputs: `data/protein_pairs/*.parquet`, each file is an aggregation of alignments for a number of taxa pairs with protein pids, source organism taxids, and alignment metrics

- Metrics: `protein_align_X`, where X is "emissions", "hits", "time", "return". The resources and used and return on investment of protein alignment.

9. **`s1.4_make_database.py`**
   Collect processed data files into a relational duckdb database.
   - Inputs: `data/taxa.parquet`, `data/taxa_thermophile_labels.parquet`, `data/protein_pairs/*.parquet`, `data/proteins/*.parquet`, `data/uniprot/proteome_metadata.csv`
   - Outputs: `data/database.ddb` relational duckdb database of taxa, proteins, taxa pairs, and protein pairs

#### Data validation

10. **`s2.1_get_hait_pairs.py`**
    Parse the protein pairs from Hait et al's excel files, query the PDB ids to get sequences.
    - Outputs: `data/validation/hait_pairs.csv`, columns include meso PDB id, thermo PDB id, and sequences
11. **`s2.2_compare_to_Tm.py`**
    Compare melting temperatures from FireProtDB and Meltome Atlas to OGTs in the dataset.
    - Inputs: `data/database.ddb`
    - Metrics: `Tm_OGT_spearman_p`, `Tm_OGT_spearman_r` Spearman R and null probability of relationship between OGT from our data and melting temperature from 3rd party dataset
12. **`s2.3_run_hait_alignment.py`**
    Compute the alignment metrics for Hait et al. pairs using identical parameters as the alignment metrics for the full dataset.
    - Params: `method` local aligner type, `method_X_params` where X is eg. "blast" params given to aligner, `blast_metrics` alignment metrics to record.
    - Inputs: `data/validation/hait_pairs.csv`
    - Outputs: `data/validation/hait_aligned_scores.csv`, columns include protein ids and alignment metrics
13. **`s2.4_compare_hait_alignment.py`**
    Compare metrics for BLAST alignments of Hait et al. pairs to our dataset, and make some plots.
    - Inputs: `data/validation/hait_aligned_scores.csv`, `data/database.ddb`
    - Outputs: `data/validation/hait_alignment/*.png` Distribution comparison of alignment metrics, and size of your dataset if we were to use Hait scores as a filter.
14. **`s2.5_get_HMM_profiles.py`**
    Download Pfam HMMs.
    - Outputs: `./data/validation/hmmer/Pfam-A.hmm`
    - Metrics: `HMM_pulled_date` date of Pfam aquisition
15. **`s2.6_hmmer_hait.py`**
    Run Pfam against proteins in Hait et al. pairs, compute Jaccard of annotations.
    - Params: `e_value` Maximum e-value for hmmer to report an annotation, `njobs` cores for parallel
    - Inputs: `data/validation/hait_pairs.csv`, `./data/validation/hmmer/Pfam-A.hmm`
    - Outputs: `data/validation/hait_scores.csv`, columns include jaccard score of Pfam annotations
    - Metrics: `mean_jaccard` Mean jaccard score of annotations over Hait pairs, `fraction_found` Fraction of proteins with at least one Pfam annotation
16. **`s2.7_run_hmmer.py`**
    Scan Pfam against all proteins on our database that are in protein pairs.
    - Params: `e_value` Maximum e-value for hmmer to report an annotation, `njobs` cores for parallel, `scan` boolean to hmmscan or hmmsearch, `prefetch` whether to

prefetch HMMs into memory or leave as file iterator, `chunk_size` size of protein chunks to run and save to file at one time.
- Inputs: `data/database.ddb`, `./data/validation/hmmer/Pfam-A.hmm`
- Outputs: `data/validation/hmmer_outputs/*.parquet` Pfam annotations for each protein, columns include the PID and a 'accessions', a semicolon separated string of Pfam accessions annotated for the PID
- Metrics: `n_proteins_in_pairs` number of proteins in pairs, `n_proteins_labeled` number of proteins with at least one Pfam annotation.

17. **`s2.8_parse_hmmer_result.py`**
Parse hmmer results into a table of protein pairs and their Jaccard scores of annotations
- Inputs: `data/validation/hmmer_outputs/*.parquet`
- Outputs: `data/validation/hmmer_labels/*.parquet`, columns include meso PID, thermo PID, and Jaccard overlap of Pfam annotations
- Metrics: `mean_jaccard` Mean jaccard score of annotations over protein pairs, `fraction_found` Fraction of proteins with at least one Pfam annotation

18. **`s2.9_compare_hait_hmmer.py`**
Compare Pfam annotations of Hait et al. pairs to our dataset, and make some plots.
- Inputs: `data/validation/hmmer_labels/*.parquet`, `data/database.ddb`
- Outputs: `data/validation/hmmer/compare_jaccard_hist.png` Distribution comparison of alignment Jaccard scores for Hait et al. pairs and our pairs.
- Metrics: `t_pvalue_base` p-value of t-test of Jaccard scores between Hait et al. and our pairs, `t_pvalue_95` p-value of t-test of Jaccard scores between Hait et al. and our pairs with > 95% blast coverage

19. **`s2.10_sample_data_for_structure.py`**
Sample some protein pairs to conduct structural alignment on, uniform over BLAST coverage.
- Params: `sample_size` number of protein pairs to sample, `metrics` list of queries to make from pairs table to sample pairs uniformly over
- Inputs: `data/database.ddb`
- Outputs: `data/validation/structure/sample_l2t_data.csv`, a sample of pairs from the pairs table, columns include meso and thermo PID, and metrics specified in params

20. **`s2.11_structure_hait.py`**
Run FATCAT structural alignment for Hait et al. pairs by getting PDB structures.
- Inputs: `data/validation/hait_pairs.csv`
- Outputs: `data/validation/structure/hait_fatcat.csv`, column include the P-value from FATCAT for the alignment

21. **`s2.12_structure_l2t.py`**
Run FATCAT structural alignment for L2T pairs by getting PDB or AlphaFold structures.
- Inputs: `data/validation/structure/sample_l2t_data.csv`
- Outputs: `data/validation/structure/l2t_sample_fatcat.csv`, column includes meso and thermo PIDs, metrics originally specified to sample uniformly from in s2.10

### Parameters
The full list of parameters and their current values for this work is given in Table S1 below:

Table S1: All parameters in the pipeline to modulate the final result.

| Stage | Parameter name | Description | Published Value |
|---|---|---|---|
| get_raw_data | min_16s_len | Minimum number of bases in 16s | 1300 |

| | | | |
|---|---|---|---|
| _taxa | | rRNA sequence to keep the sequence | |
| | max_16s_len | Maximum number of bases in 16s rRNA sequence to keep the sequence | 1600 |
| parse_proteins | max_prot_per_file | Number of proteins per chunk for saving UniProtKB as minimal table | 100000 |
| label_taxa | ogt_threshold | Binary threshold to consider a taxa thermophilic, not inclusive | 40.0 |
| get_16s_blast_scores | num_threads | BLAST cpus for computation | 20 |
| | word_size | Size of initial exact nucleotide matching in BLAST | 28 |
| | gapopen_penalty | Penalty for bit score incurred by starting a gap in the alignment | 2 |
| | gapextend_penalty | Penalty for bit score incurred by extending a gap in the alignment | 1 |
| | reward | Score increase for a match in alignment | 1 |
| | penalty | Penalty for mismatch in alignment | -2 |
| | ungapped | Whether to do an alignment without allowing gaps | false |
| | blast_metrics | List of metrics defined in learn2therm.blast.BlastMetrics to compute for 16s rRNA alignment | * See note 1 below |
| label_all_pairs | blast_metric_thresholds | Nested dictionary of name of alignment metric and two fields "thresh" which is the threshold for considering a pair, non inclusive, and "greater", a boolean of whether we want bigger or smaller than the threshold | * See note 2 below |
| get_protein_blast_scores | dask_cluster_class | Class from Dask Jobqueue[1] to use for parallel workers | SLURMCluster |
| | max_protein_length | Maximum number of amino acids, inclusive, to keep for protein pair search | |

| | | | |
|---|---|---|---|
| | method | Which aligner to use, one of ‘blast’ or ‘diamond’ | diamond |
| | n_jobs | Number of independent parallel workers to keep running | 80 |
| | save_frequency | Number of taxa pairs between DVC checkpointing | 20000 |
| | method_blast_params | Parameters for BLASTp algorithm | See Table S2 below. |
| | method_diamond_params | Parameters for DIAMOND algorithm | See Table S2 below. |
| | blast_metrics | List of metrics defined in learn2therm.blast.BlastMetrics to compute for protein alignment | * See note 1 below |
| run_hmmer | e_value | Maximum E value for labelling Pfam annotations with HMMER | 1.e-10 |
| | chunk_size | Number of vector chunks to run at a time | 2000 |
| | prefetch | Boolean of whether to load HMMs into memory, or leave as disk iterator | true |
| | njobs | Number of CPUs to use for parallel pyhmmer | 32 |
| | scan | Whether to use HMMScan, alternative HMMSearch | false |
| sample_data_for_structure | sample_size | Number of protein pairs to keep for structural alignment | 10,000 |
| | metrics | Queries of protein pairs table to compute and use for sampling, the columns is binned into 5 bins and samples uniformly from those bins | [“(query_align_cov+subject_align_cov)/2.0"] |

Table S2: Parameters for BLASTp, if using it for protein alignment

| **Parameter name** | **Description** | **Published Value** |
|---|---|---|
| num_threads | CPUs for each worker to run BLASTp | 6 |
| word_size | Size of initial Amino Acid matching in BLASTp | 3 |
| gapopen | Penalty to bit score for opening a gap in alignment | 11 |

| | | |
|---|---|---|
| gapextend | Penalty to bit score for extending a gap | 1 |
| matrix | Substitution matrix for comparing amino acids in alignment | BLOSUM62 |
| threshold | Minimum score for word to be added to lookup table | 11 |
| ungapped | Whether to run alignment without gaps | false |
| evalue | Maximum E value to keep scoring alignment | 0.00001 |
| qcov_hsp_perc | Minimum percent coverage of query strand to keep alignment | 75 |

Table S3: Parameters for DIAMOND, if using it for protein alignment

| **Parameter name** | **Description** | **Published Value** |
|---|---|---|
| num_threads | CPUs for each worker to run DIAMOND | 6 |
| sensitivity | How sensitive the search should be for initial matching | ultra-sensitiv e |
| gapopen | Penalty to bit score for opening a gap in alignment | 11 |
| gapextend | Penalty to bit score for extending a gap | 1 |
| matrix | Substitution matrix for comparing amino acids in alignment | BLOSUM62 |
| iterate | Whether to start with lower sensitivity alignment | false |
| global_ranking | Limit on the number of Smith Waterman extensions per query, with the target sequences ranked by their ungapped extension scores. | null |
| evalue | Maximum E value to keep scoring alignment | 0.00001 |
| hsp_cov | Minimum percent coverage of query and subject strand to keep alignment | 75 |

* *Note 1:* Any metric that is a method name of the class learn2therm.blast.BlastMetrics can be specified here. See Section S2 for definitions of these metrics. The values used in this work are: [ local_gap_compressed_percent_id, scaled_local_query_percent_id, scaled_local_symmetric_percent_id, local_E_value, query_align_start, query_align_end, subject_align_end, subject_align_start, query_align_len, query_align_cov, subject_align_len, subject_align_cov, bit_score ]

* *Note 2:* To threshold by E value less than $1e^{-10}$, we would have a nested specification like the following:

blast_metric_thresholds:
 local_E_value:
  thresh: 1e-10
  greater: false

### S2. Alignment metrics

A number of metrics were computed for pairwise alignments, and are given in the database. Their definition, description, and database tag is provided below.

Given an alignment of length L including gaps upon query sequence and subject sequence with lengths A and B, a column at position $i \in$ [0,L] has:

- $I_i = 1$ if the column is a match between the two strands, 0 otherwise
- $G_i = 1$ if the column has a gap in the alignment, 0 otherwise
- $G_i^X = 1$ if the column has a gap in the X strand, 0 otherwise
- $GF_i = 1$ if the column has a gap in the alignment and has no gap column to its left, 0 otherwise
- $S_i$, the bit score of the alignment according to a substitution matrix (BLOSUM62 was used, however this is a free parameter)

$N(X)$ is used to notate $\sum_{i=0}^{L} X_i$.
$M$ is the number of sequences scanned for alignments.

Table S4: Names and definitions of alignment metrics computed.

| Tag | Description | Definition |
|---|---|---|
| local_gap_compressed_percent_id | Percent identity of the alignment, with contiguous gaps counted as only one. | $\frac{N(I)}{L - N(G)+N(GF)}$ |
| scaled_local_query_percent_id | Percent identity normalised to the query strand | $\frac{N(I)}{A}$ |
| scaled_local_symmetric_percent_id | Percent identity normalised to the average strand length | $\frac{2\,N(I)}{A+B}$ |
| bit_score | BLAST bit score | $N(S)$ |
| local_E_value | BLAST E value | $\frac{M\,A}{2^{N(S)}}$ |
| query/subject_align_cov | Coverage of alignment on strand | $\frac{L - N(G^{query})}{A}, \frac{L - N(G^{subject})}{B}$ |

Note: Start and end positions of the alignment on a strand is also tracked as query/subject_align_start/end. The difference between start and end, e.g., the length of the alignment per strand is tracked as query/subject_align_len.

### *S3. 16s rRNA Alignment*

Taxa Pairs were determined via alignment of 16s rRNA strands from Refseq using BLASTn 2.14.0+. The parameters for the search are shown in Table S5 below, default if not listed:

Table S5: BLASTn parameters used.

| Parameter | Value |
|---|---|
| word_size | 28 |
| gapopen | 2 |
| gapextend | 1 |
| reward | 1 |
| penalty | -2 |
| ungapped | false |
| evalue | 1.0 |
| perc_identity | 0.5 |

The best HSP as determined as the one with the most coverage of both strands (averaged) was then retained for each putative pair. Taxa were labelled as pairs if the resulting coverage for both strands was >=98.5% and the gap compressed percent identity >=81%.

### *S4. Protein Alignment*

#### Reported alignment

Protein Pairs were determined via alignment of UniProtKB sequence using DIAMOND version 2.1.6, a BLAST-like software. The parameters for the search are shown in Table S6 below, default if not listed:

Table S6: DIAMOND parameters used.

| Parameter | Value |
|---|---|
| word_size | 3 |
| sensitivity | ultra-sensitive |
| iterate | false |
| matrix | BLOSUM62 |
| global_ranking | null |
| gapopen | 11 |
| gapextend | 1 |
| evalue | 0.00001 |

| | |
|---|---|
| max_hsps | 100 |
| id | 0 |
| query-cover | 75 |
| subject-cover | 75 |

The best HSP as determined as the one with the most coverage of both strands (averaged) was then retained for the pair and is reported in the dataset with a number of alignment metrics (see Section S2).

### Cost of alignment

Before running protein alignment on many taxa pairs and potentially wasting compute, we compared the use of BLASTp to DIAMOND using identical parameters wherever possible, eg. word size, matrix, penalties, etc. This was conducted for 10k random proteins vs 10k random proteins. The compute time, carbon, and return on investment in terms of number of hits is shown below if Figure S1. We chose to use DIAMOND for alignment given the small decrease in sensitivity compared to near order of magnitude cost reduction.

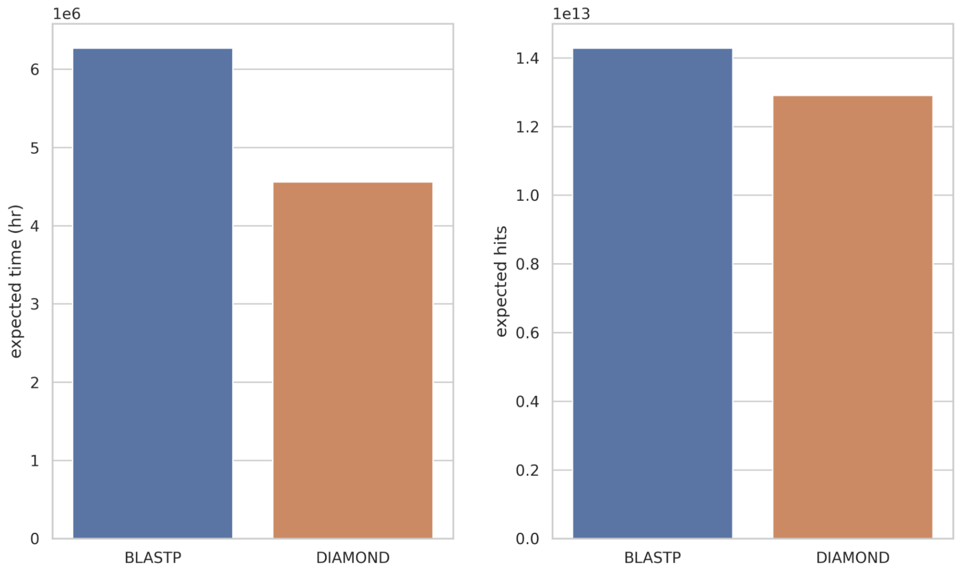

Figure S1: BLASTp vs DIAMOND resource test of protein alignment. DIAMOND significantly reduces cost while retaining most of the sensitivity.

### *S5. Data Schema*

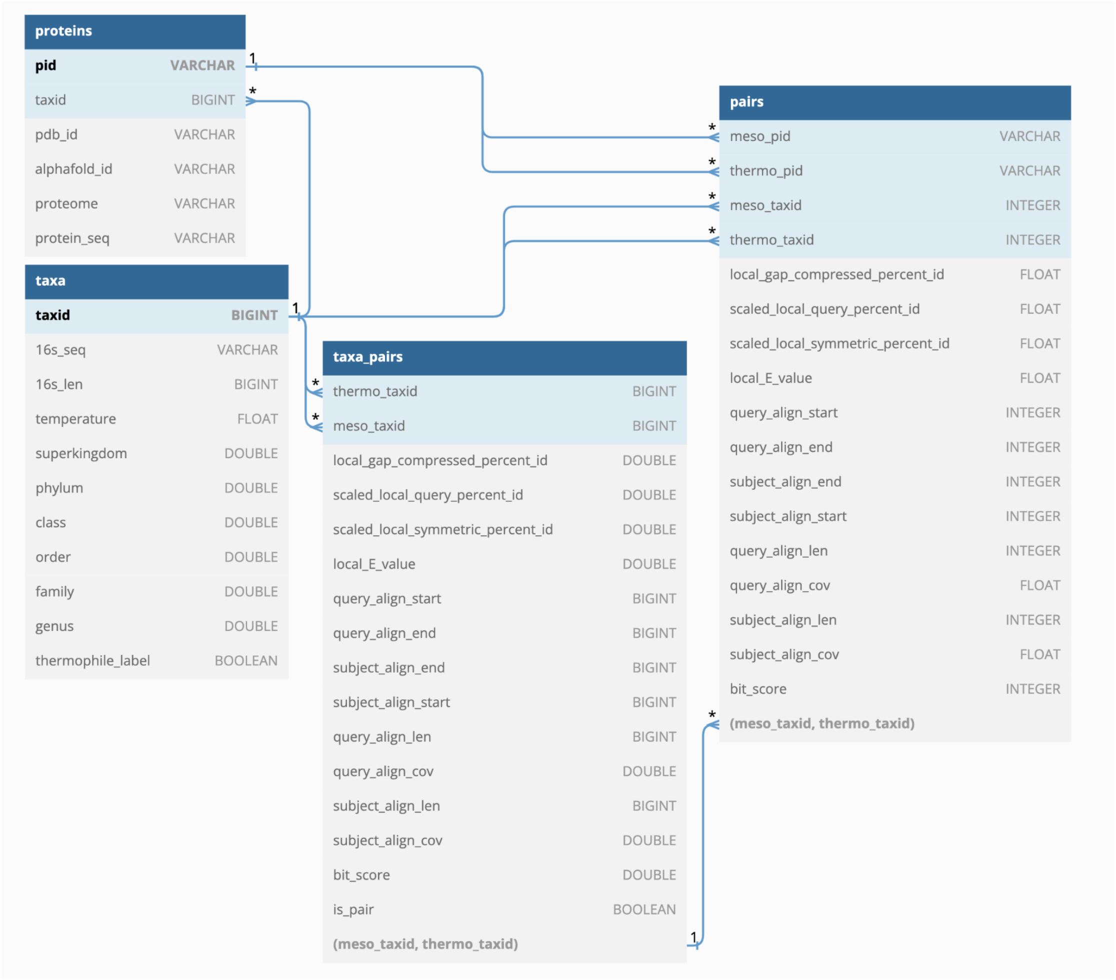

Figure S2: Schema for presented database. The 'taxa' and 'proteins' tables contain raw data from UniProtKB, RefSeq, and Engqvist. The 'taxa_pairs' and 'pairs' tables contain information on alignment of 16s rRNA and protein sequences. See Table S7 below for a description of each field in the database.

Table S7: Data fields within the presented relational database.

| Table | Field | Description |
|---|---|---|
| taxa | taxid | Primary key, NCBI taxid |
| taxa | 16s_seq | Nucleotides of 16s rRNA sequence |
| taxa | 16s_len | Length of 16s rRNA strand |
| taxa | temperature | Optimal Growth Temperature |
| taxa | superkingdom | NCBI superkingdom integer identifier |
| taxa | phylum, …, genus | NCBI integer identifier for taxonomy |
| taxa | thermophile_label | Indicator if organism is a thermophile as determined by pipeline parameters |

| | | |
|---|---|---|
| proteins | pid | Primary key, UniProtKB identifier |
| proteins | taxid | NCBI taxid of host organism, foreign key to taxa |
| proteins | pdb_id | PDB cross reference if present |
| proteins | alphafold_id | AlphaFold database cross reference |
| proteins | proteome | Identifier of UniProtKB proteome that protein is a part of, if present |
| proteins | protein_seq | Amino acids of the sequence |
| taxa_pairs | thermo_taxid | NCBI taxid of thermophile, foreign key to taxa. Serves as compound primary key with meso_taxid |
| taxa_pairs | meso_taxid | NCBI taxid of mesophile, foreign key to taxa. Serves as compound primary key with thermo_taxid |
| taxa_pairs | is_pair | Indicator if pair of taxa is considered a pair by pipeline parameters for downstream protein alignments. |
| taxa_pairs | * | A number of alignment metrics, see section S2 |
| pairs | meso_pid | UniProtKB identifier of mesophilic protein, foreign key to proteins. Serves as compound primary key with thermo_pid |
| pairs | thermo_pid | UniProtKB identifier of thermophilic protein, foreign key to proteins. Serves as compound primary key with meso_pid |
| pairs | meso_taxid | NCBI taxid of mesophile, foreign key to taxa. Serves as compound foreign key with thermo_taxid to taxa_pairs |
| pairs | thermo_taxid | NCBI taxid of thermophile, foreign key to taxa. Serves as compound foreign key with meso_taxid to taxa_pairs |
| pairs | * | A number of alignment metrics, see section S2 |

### *S6. Pfam annotations*

Pfam was used to annotate proteins in protein pairs. For a given protein pair, both proteins were run against 19,632 HMMs from Pfam version 35.0 using pyhmmer.[2] All annotations with sequence hit E-value < $1e^{-10}$ were retained. The set Jaccard score was then computed for annotations between two proteins, as defined below:

$$J = \frac{\cap(A, B)}{\cup(A, B)} \quad (Eqn\ 1)$$

Where $A$, and $B$ are the set of Pfam annotations from Protein A and B, respectively. Thus, if protein A is labelled with families "pfam_0" and "pfam_1", while protein B is labelled with only "pfam_0", the Jaccard score is 0.5. If Pfam was unable to find annotations for one of two proteins within a pair meeting the minimum HMMER requirements, the Jaccard score is 0.0. Note that protein pairs where neither protein was annotated were ignored for analysis, likely falling into an unknown protein family.

### S7. Deep Predictor Training

We fine-tuned ProtBERT on proteins from our dataset to act as a classifier of thermophilic or mesophilic from sequence alone.[3] To prepare for this fine-tuning, we preprocessed the data as follows.

The N=10 mil proteins with sequence length less than or equal to 250 amino acids from our database was considered. Data labelled with OGT in [30°C, 60°C) were excluded. Proteins with higher temperature were labelled thermophilic while the others were labelled mesophilic. The resulting data was randomly downsampled in order to produce a balanced dataset of 175k each of thermophilic and mesophilic proteins. In order to minimise redundancy in the dataset, we conducted MinHash distance removal of sequence tokens, a common step for reducing leakage in traditional natural language datasets.[4–6] Here, we compute the set of all 3-mers for each amino acid sequence, then computed the MinHash of the resulting set using 100 permutations. Proteins were clustered using distance of the MinHash Jaccard score > 0.6, and only a single protein was retained from each group. The dataset was finally split into development and test set with a fraction of 10% test using random splitting on the protein's taxid, thus no two proteins from the same organism occur across datasets.

The model itself was downloaded with pre-trained weights from Huggingface, and a new MLP layer was created to act as a binary predictor. Specifically, the pretrained embeddings of size 1024 from the output of ProtBERT were pooled using a linear projection of size 1024 and then mean over amino acids in the sequence, followed by a two-neuron binary predictor. The pytorch readout for the model is given below, note that neural modules are from Huggingface's ecosystem. See their model classes here.[7]

```
BertForSequenceClassification(
 (bert): BertModel(
   (embeddings): BertEmbeddings(
     (word_embeddings): Embedding(30, 1024, padding_idx=0)
     (position_embeddings): Embedding(40000, 1024)
     (token_type_embeddings): Embedding(2, 1024)
     (LayerNorm): LayerNorm((1024,), eps=1e-12, elementwise_affine=True)
     (dropout): Dropout(p=0.0, inplace=False)
   )
   (encoder): BertEncoder(
     (layer): ModuleList(
       (0-29): 30 x BertLayer(
         (attention): BertAttention(
           (self): BertSelfAttention(
             (query): Linear(in_features=1024, out_features=1024, bias=True)
             (key): Linear(in_features=1024, out_features=1024, bias=True)
             (value): Linear(in_features=1024, out_features=1024, bias=True)
```

```
            (dropout): Dropout(p=0.0, inplace=False)
          )
          (output): BertSelfOutput(
            (dense): Linear(in_features=1024, out_features=1024, bias=True)
                                  (LayerNorm):   LayerNorm((1024,),   eps=1e-12,
elementwise_affine=True)
            (dropout): Dropout(p=0.0, inplace=False)
          )
        )
        (intermediate): BertIntermediate(
          (dense): Linear(in_features=1024, out_features=4096, bias=True)
          (intermediate_act_fn): GELUActivation()
        )
        (output): BertOutput(
          (dense): Linear(in_features=4096, out_features=1024, bias=True)
                                  (LayerNorm):   LayerNorm((1024,),   eps=1e-12,
elementwise_affine=True)
          (dropout): Dropout(p=0.0, inplace=False)
        )
      )
    )
  )
  (pooler): BertPooler(
    (dense): Linear(in_features=1024, out_features=1024, bias=True)
    (activation): Tanh()
  )
 )
 (dropout): Dropout(p=0.0, inplace=False)
 (classifier): Linear(in_features=1024, out_features=2, bias=True)
)
```

The model was then fine tuned for two epochs on the 280k training proteins. The training parameters are given below:

```
TrainingArguments(
_n_gpu=1,
adafactor=False,
adam_beta1=0.9,
adam_beta2=0.999,
adam_epsilon=1e-08,
auto_find_batch_size=False,
bf16=False,
bf16_full_eval=False,
data_seed=None,
dataloader_drop_last=False,
dataloader_num_workers=0,
```

```
dataloader_pin_memory=True,
ddp_bucket_cap_mb=None,
ddp_find_unused_parameters=None,
ddp_timeout=1800,
debug=[],
deepspeed=None,
disable_tqdm=False,
do_eval=True,
do_predict=False,
do_train=True,
eval_accumulation_steps=25,
eval_delay=0,
eval_steps=6,
evaluation_strategy=steps,
fp16=True,
fp16_backend=auto,
fp16_full_eval=False,
fp16_opt_level=O1,
fsdp=[],
fsdp_min_num_params=0,
fsdp_transformer_layer_cls_to_wrap=None,
full_determinism=False,
gradient_accumulation_steps=25,
gradient_checkpointing=True,
greater_is_better=False,
group_by_length=False,
half_precision_backend=cuda_amp,
hub_model_id=None,
hub_private_repo=False,
hub_strategy=every_save,
hub_token=<HUB_TOKEN>,
ignore_data_skip=False,
include_inputs_for_metrics=False,
jit_mode_eval=False,
label_names=None,
label_smoothing_factor=0.0,
learning_rate=5e-05,
length_column_name=length,
load_best_model_at_end=True,
local_rank=0,
log_level=info,
log_level_replica=passive,
log_on_each_node=True,
logging_dir=./data/ogt_protein_classifier/model/runs/Jun19_12-16-35_g3070,
logging_first_step=False,
logging_nan_inf_filter=True,
```

```
logging_steps=1,
logging_strategy=steps,
lr_scheduler_type=linear,
max_grad_norm=1.0,
max_steps=-1,
metric_for_best_model=loss,
mp_parameters=,
no_cuda=False,
num_train_epochs=2,
optim=adamw_hf,
optim_args=None,
output_dir=./data/ogt_protein_classifier/model,
overwrite_output_dir=False,
past_index=-1,
per_device_eval_batch_size=32,
per_device_train_batch_size=32,
prediction_loss_only=False,
push_to_hub=False,
push_to_hub_model_id=None,
push_to_hub_organization=None,
push_to_hub_token=<PUSH_TO_HUB_TOKEN>,
ray_scope=last,
remove_unused_columns=True,
report_to=['tensorboard', 'codecarbon'],
resume_from_checkpoint=None,
run_name=./data/ogt_protein_classifier/model,
save_on_each_node=False,
save_steps=6,
save_strategy=steps,
save_total_limit=None,
seed=42,
sharded_ddp=[],
skip_memory_metrics=True,
tf32=None,
torch_compile=False,
torch_compile_backend=None,
torch_compile_mode=None,
torchdynamo=None,
tpu_metrics_debug=False,
tpu_num_cores=None,
use_ipex=False,
use_legacy_prediction_loop=False,
use_mps_device=False,
warmup_ratio=0.0,
warmup_steps=0,
weight_decay=0.0,
```

```
xpu_backend=None,
)
```

The final model model was saved and evaluated against the test set. It can be found on the Huggingface Hub, here.[8]

### *S8. ESM Mapping*

To produce a 2D mapping of proteins (Figure 1B), we used ESM2 embeddings.[9] First, embeddings from ESM Atlas were retrieved for all proteins with Tm>0.9 and pLDDT>0.9. These are the proteins that the ESM language model's structural prediction module is most confident in, eg. those that are not likely extrapolative given the training sequences/structures. The embeddings represent the mean over sequence tokens of the pretrained model's output latent space, producing a single size 2,560 vector for each protein sequence.

Due to computational cost, we sampled data in order to make dimensionality reduction achievable. Only 500k proteins from the ESM Atlas were retained by random sampling. We then randomly sampled proteins from pairs from ours and Hait et al.'s dataset, retaining the ratio of dataset sizes to each other and to the ESM Atlas. This resulted in 240k proteins from our dataset, and 82 from Hait et al.'s. We then used the same pretrained model version ('`esm2_t36_3B_UR50D`') as was used for ESM Atlas to embed our's and Hait et al.'s proteins.

Finally, we ran multicore T-SNE on the embedded vectors down to two dimensions.[] We used the following parameters, the rest default:

Table S8: Parameters used for T-SNE dimensionality reduction of ESM embeddings.

| **Parameter** | **Value** |
|---|---|
| n_iter_early_exag | 500 |
| n_iter | 2000 |
| perplexity | 30.0 |
| theta | 0.5 |

### *References*

1. Dask-Jobqueue — Dask-jobqueue 0.8.2+0.gff47d71.dirty documentation. https://jobqueue.dask.org/en/latest/.
2. Larralde, M. & Zeller, G. PyHMMER: a Python library binding to HMMER for efficient sequence analysis. *Bioinformatics* **39**, btad214 (2023).
3. Elnaggar, A. *et al.* ProtTrans: Towards Cracking the Language of Life's Code Through Self-Supervised Learning. 2020.07.12.199554 Preprint at https://doi.org/10.1101/2020.07.12.199554 (2021).
4. High-Dimensional Similarity Query Processing for Data Science | Proceedings of the 27th ACM SIGKDD Conference on Knowledge Discovery & Data Mining. https://dl.acm.org/doi/abs/10.1145/3447548.3470811.
5. Team, T. A. Text Similarity using K-Shingling, Minhashing and LSH(Locality… – Towards AI. https://towardsai.net/p/l/text-similarity-using-k-shingling-minhashing-and-lshlocality-sensitive-hashing, https://towardsai.net/p/l/text-similarity-using-k-shingling-minhashing-and-lshlocality-sensitive-hashing.
6. Qurashi, A. W., Holmes, V. & Johnson, A. P. Document Processing: Methods for Semantic Text Similarity Analysis. in *2020 International Conference on INnovations in Intelligent SysTems and Applications (INISTA)* 1–6 (2020). doi:10.1109/INISTA49547.2020.9194665.
7. Huggingface. BERT. https://huggingface.co/docs/transformers/v4.30.0/en/model_doc/bert.
8. Komp, Evan & Beck, David. learn2therm_model. *Huggingface* doi:10.57967/hf/0815.
9. Choudhuri, S. Protein Folding Problem: A Comparative Analysis of ESM2. SSRN Scholarly Paper at https://doi.org/10.2139/ssrn.4301467 (2022).
